# Dissecting the sources of variation in neuronally differentiated iPSC lines through multi-omics analysis

**DOI:** 10.64898/2026.06.10.731279

**Authors:** Casper de Visser, Lisa Rahm, Elly Lewerissa, Rachel Mijdam, Cenna Doornbos, Junda Huang, Luke O’Gorman, Firdaws Badmus, Clara D.M. van Karnebeek, Catharina G. Faber, Judith Verhoeven, Hans van Bokhoven, Nael Nadif Kasri, Dirk J. Lefeber, Peter A.C. ’t Hoen, Alain J. van Gool, Purva Kulkarni

## Abstract

Induced pluripotent stem cells (iPSCs) are widely used as patient-specific disease models, yet substantial unexplained variability in molecular and functional readouts limits their reliability. Here, we systematically investigated the sources of variation in iPSC-derived neurons for three rare genetic disorders: Myotonic Dystrophy Type 1, chromodomain-DNA-helicase-binding protein 2-related disorder and *N*-acetylneuraminic acid synthase deficiency. This was performed by profiling multi-omics layers: genomics, epigenomics, transcriptomics, proteomics, metabolomics and lipidomics. Our study found that clonal variability was comparable to inter-patient differences and that neuronal differentiation state and nutrient-driven metabolic activity emerged as dominant contributors to variability observed across omics layers. Clonal differences could partly be attributed to stochastic differences in DNA methylation established during reprogramming. By modeling and correcting the observed variation, we improved the detection of disease-associated molecular signatures. Our study provides guidelines for improved study design and data analysis to minimize variability, enabling robust biomarker discovery and reliable iPSC-based disease modeling.

## Introduction

Induced pluripotent stem cells (iPSCs) are powerful *in vitro* cellular models to are frequently used to study disease mechanisms and interventions [1–5]. They represent a powerful platform for mechanistic and translational research, as they can be generated directly from patient tissues while retaining the individual’s genetic background [6,7]. Additionally, iPSCs can reflect disease-specific phenotypes, as they can be derived from patients themselves and can be differentiated into specific cell types relevant to the disease of interest. For example, iPSCs can be differentiated into neurons to enable the modelling of neurological disorders [8]. These characteristics make iPSC models a potentially powerful and a more ethically accepted alternative to traditional animal models.

Previous studies [6,9–14] highlighted considerable variability in iPSC omics measurements, including RNA, protein, and metabolite levels. Although most of the variation was reported as donor-specific [6], additional variation was discovered to be influenced by non-genetic factors. For example, the reprogramming of background cells to iPSC was shown to affect DNA methylation [13]. Other studies utilizing transcriptomics and DNA methylation data analysis demonstrated clonal heterogeneity that affected the cardiac differentiation potential of iPSCs [14]. This variability can affect experimental outcomes and thus become a challenge for the consistency and reproducibility of iPSC-based research. These issues are particularly pressing when robust, standardized readouts are essential, for example when validating models across laboratories for drug response- or during regulatory evaluation of therapy development. Altogether, it is of essence to reveal the sources of variation and use this information to improve the reproducibility of iPSC-based studies.

Here, we focus on uncovering key sources of variation in iPSC models of three neurological diseases through multi-omics analysis to support the development of reliable disease models. Specifically, we characterized these variations by analyzing biomolecular measurements across multiple omics layers: genomics, epigenomics, transcriptomics, proteomics, metabolomics, lipidomics. We investigated multiple samples from patients diagnosed with Myotonic Dystrophy Type 1 (DM1) [15,16], chromodomain-DNA-helicase-binding protein 2 (CHD2) related disorder [17], and *N*-acetyl-d-neuraminic acid synthase congenital disorders of glycosylation (NANS-CDG) [18] (Fig 1).

**Fig 1.**
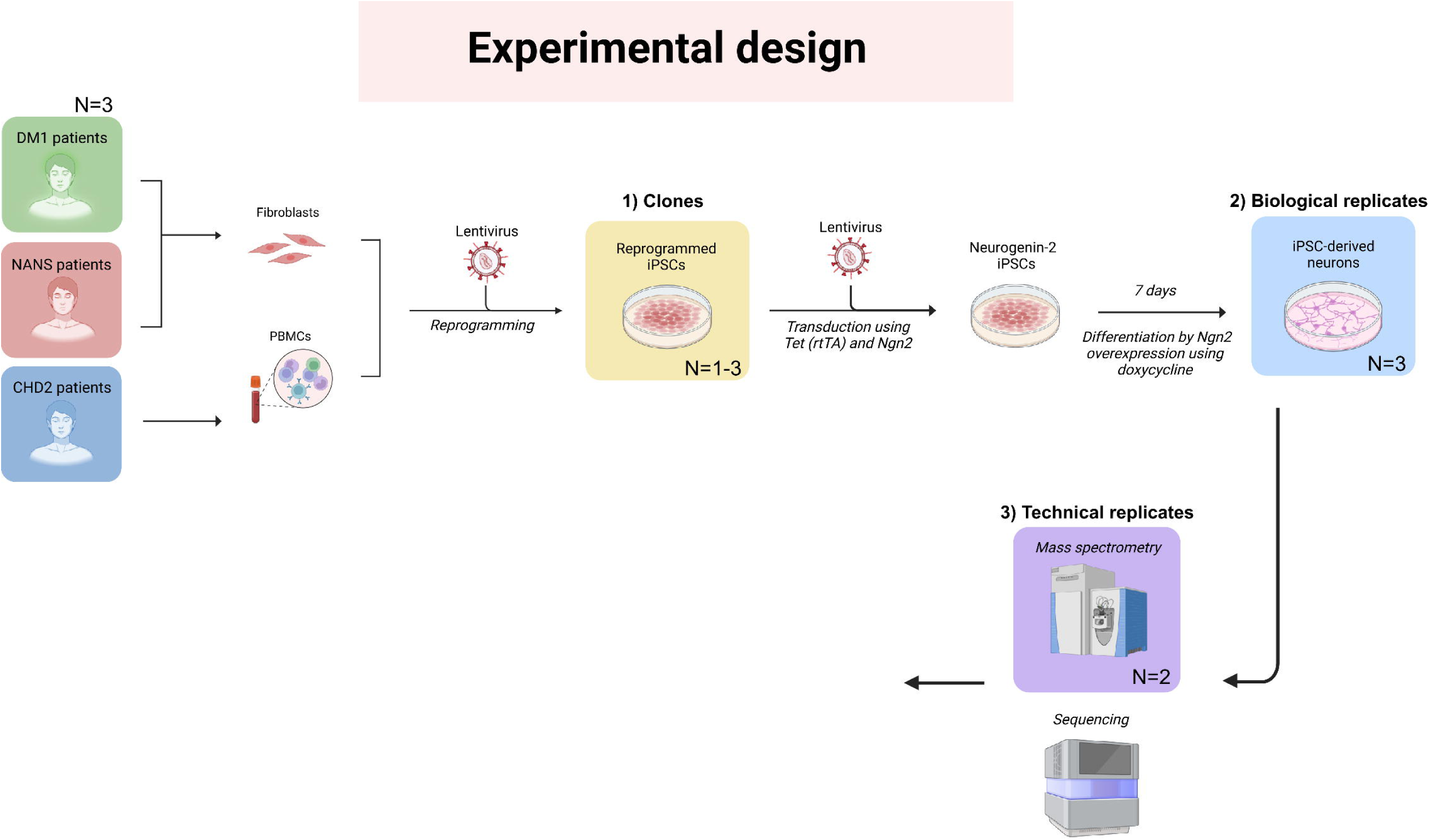
Study design. iPSC-derived neurons were generated for three patients per disease group, and control lines (Fig S1). Fibroblasts were used as starting tissue for all cell lines, except for the CHD2 cell lines which were created from PBMCs. Reprogrammed iPSCs were differentiated 7 days *in vitro* by *Ngn2* overexpression induced by doxycycline. This process was repeated three times for each cell line, which we refer to as biological replicates. The resulting cell lines were used for multi-omics profiling. The total number of samples analyzed in this study is 44. Missing samples in omics technology are greyed out. The total number of omics features in each omics layer is also indicated.

## Results

### Study design

Our study compared omics datasets generated from cell cultures derived from different patient groups. For each disease, we selected cell lines from three patient donors (two females and one male) along with three controls. Cell lines were derived from fibroblasts or peripheral blood mononuclear cells (PBMCs) through lentiviral reprogramming into iPSCs (Supplementary Table 1, Fig 1). In addition, we generated two isogenic lines using CRISPR-Cas9: a CHD2-mutant line created by editing the control line 409B, and a corrected DM1 control line generated by removing the pathogenic repeat expansion from DM1 patient line 1 (now referred to as DM1-P1) (Fig S1).

To systematically capture sources of variation, we defined three levels of replicates (Fig 1). For some patients, multiple independent iPSC lines were used, which will be referred to as clones. Moreover, each iPSC line was differentiated into neurons in 3 independent experiments and cultured for 7 days *in vitro* by the same experimenter. These differentiation replicates will be referred to as the biological replicates in this study. Lastly, for specific analyses and specific omics data sets (proteomics & metabolomics) duplicate measurements on the same sample were available as well, which will be referred to as technical replicates. With this sample hierarchy, we aimed to identify the main sources of variation in iPSC-derived neurons, and to correct for these sources of unwanted variation, enhancing detection of disease-specific signals in the multi-omics data.

Our multi-omics dataset included six omics layers from 44 samples: whole exome sequencing (WES), enzymatic methylation sequencing (EMseq), RNA sequencing (RNAseq), mass spectrometry (MS)-based proteomics, and untargeted MS-based metabolomics measured at two centers namely, the Radboud university medical center (RUMC) and the Leiden University Medical Center (LUMC), MS-based targeted and untargeted lipidomics. (Fig 1). By analyzing these omics levels measured in the iPSC-derived neurons (Fig S2, Fig S3), and applying single and multi-omics analyses, we aimed to explain the origin of the dominant sources of variation.

### Sources of variability in omics measurements

We performed a range of single- and multi-omics analyses to identify the key sources of variation in our dataset from iPSC-derived neurons, as detailed in Figure S2 and Figure S3.

#### Multi-omics data quality control

Because analyzing multiple omics data sets increases the risk of sample mislabeling and cross-dataset mismatches we performed quality control (QC) using sample concordance analyses. We identified possible sample mix-ups using sex-concordance analyses (see Methods). We conclusively validated these potential sample mix-ups using an extensive single nucleotide polymorphism (SNP)-concordance analysis based on all available sequencing data (see Methods). After correction for these sample mix-ups we continued downstream analyses.

Additionally, we inspected the copy number variation (CNV)-WES data generated from the undifferentiated iPSCs. This revealed genetic variations that were present among clones of the same patient line. For example, in DM1 patient 2 clone 2 (DM1-P2-C2) we found an 84 kb deletion on chromosome 12, encompassing *NECAP1, FOXJ2, C3AR1* and *CLEC4A*. Because none of the detected CNVs were expected to influence the multi-omics data at a large scale, we proceeded with analyses using this set of iPSC lines.

#### Principal Component Analysis demonstrates disease-specific variation in the data

PCA was then applied to the individual omics data sets. For the first 10 principal components (PCs), which explain most of the variation in each data set, we checked associations with both biological (disease group, sex) and technical (tissue, culturing strategy) variables (Fig S5). For all omics layers, we associated one or multiple of the top 10 PCs with differences between disease groups, indicative of the quality of our data and their relevance to the diseases studied. This was strongest in the EMseq data, where most variation explained by the PCs was associated with differences between patient groups. This can be partially explained by the specific selection of methylation sites in this data. Due to the large size of the complete set of methylation sites (27+ million), we selected the 100,000 most variable methylation sites across all samples for this particular data set. When methylation sites were randomly selected, the joint variation explained by disease-linked PCs was reduced (Fig S5), which is expected as most CpG sites are non-variable [19].

In addition, PCA indicated that all omics layers were also affected by experimental factors, such as tissue origin and cell culturing strategy. Some of these associations between the PCs and these factors could have been confounded by disease. For example, separation of CHD2 patients from controls and other disease groups may also be confounded by tissue differences, as only CHD2 patient lines were generated from peripheral blood mononuclear cells (PBMCs).

In summary, PCA indicated that all omics layers exhibited disease-associated variation. Thus, these data sets were suitable for the discovery of disease-specific variation. However, the majority of variation is not explained by the disease, and we performed additional analyses to identify these sources of variation.

#### Pairwise correlations reveal differences among patients and clones

To assess the variation between samples, we determined the pairwise correlations of all omics data (degree of methylation for each CpG site or mRNA, protein, metabolite, lipid abundance). Owing to the sample hierarchy in our data (disease > patient > clone > replicate) (Fig 1), we were able to systematically compare correlations between closely related samples against those that were less related. For instance, we analyzed whether correlations within the same disease group differed from correlations across different disease groups.

To achieve this, we calculated correlations across unrelated samples, samples within the same disease group, and among biological and technical replicates. R-squared (R^2^) values of these correlations were visualized for each omics data set (Fig 2). As expected, technical replicates, only available for proteomics and metabolomics, displayed the highest concordance, with mean R^2^ values of 0.99 and 0.98, respectively. As expected, gradually lower correlations were observed for biological replicates, samples in the same disease group and unrelated samples (Fig 2A). For the omics datasets for which technical replicates were not available, we observed the same overall pattern and high correlations between biological replicates (Fig S6).

**Fig 2.**
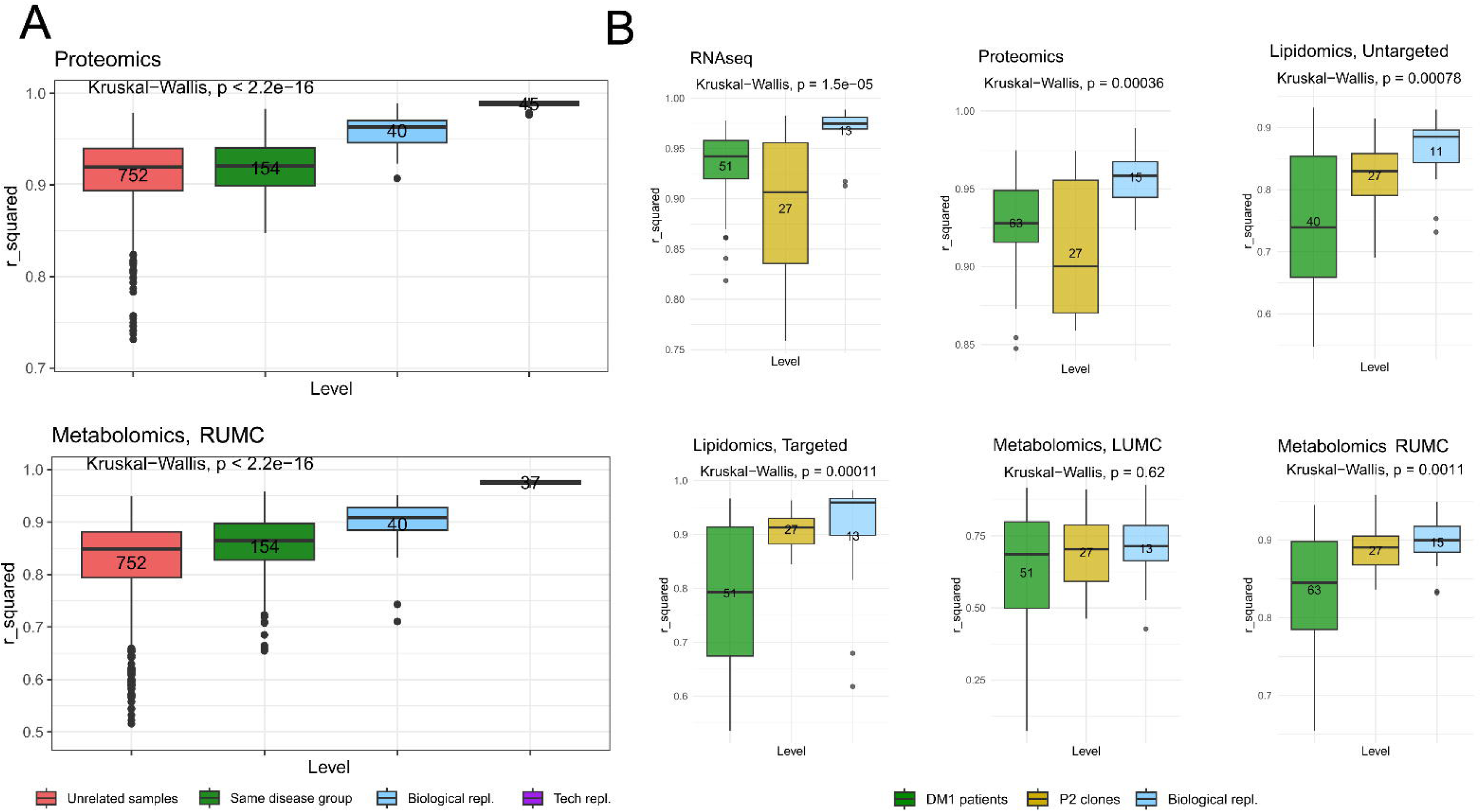
Pairwise correlations of measured omics levels. A) Pairwise correlations of metabolomics (RUMC) and proteomics levels between unrelated samples (red), samples from iPSC lines in the same disease group (green), samples from the same iPSC lines differentiated on different days (biological replicates, blue), and technical replicates (purple). Significant differences among the R^2^ values were assessed using the Kruskal-Wallis test. B) Pairwise correlations of all omics data set among DM1 patients, including correlations between samples from different DM1 patients (green), different clones from the same patient (P2) (yellow) and biological replicates (blue). Boxplots represent the median (horizontal line), boxes the interquartile ranges (IQR), whiskers represent the variability outside the upper and lower quartiles, excluding outliers that are marked as individual data points.

For some patient lines (DM1-P2 and CHD2-P3), multiple iPSC clones were available. Interestingly, correlations between different clones from the same patient were sometimes as low as those between different patients within the same disease group. For example, proteomics and transcriptomics correlations among the DM1-P2-C2 lines were on average lower (R^2^=0.90; R^2^=0.89, respectively) than among cell lines from different DM1 patients (R^2^=0.92; R^2^=0.93, respectively) (Fig 2B). When inspected in more detail, the proteomics and transcriptomics data measured in clone 2 were found to be consistently divergent from the other clones of the same patient line (Fig S7).

In contrast, correlations of the CHD2-P3 clones did not show differences as large as those for the DM1-P2 clones (Fig S8). However, correlations of the metabolomics and lipidomics data were on average lower (0.8) for the CHD2 patient lines, also for biological replicates, when comparing to the other pairwise correlations of these omics data (Fig 2A).

In conclusion, the pairwise correlations demonstrated clear clonal differences, which asked for further characterization in subsequent analyses.

#### Differences in expression of neuronal markers reflect variability in neuronal differentiation

To assess neuronal differentiation status of the iPSC-derived neurons after 7 days of differentiation, we examined gene and protein expression levels of both early and mature neuronal markers, and pluripotency markers.

The transcriptomics data were summarized in a heatmap and samples were clustered based on these markers (Fig 3). Consistent with the pairwise correlations, the DM1-P2-C2 stood out from the other clones. Both early (*TUBB3, MAP2, TBR1, NEUROG2, SATB2*) and mature (*RBFOX3, NEFL, NEFM, SYN1, SYP, NEFH*) neuronal markers were consistently lower expressed in this clone, with the strongest reduction observed in the mature markers. In line with these findings, pluripotency markers were expressed higher in the DM1-P2-C2 samples compared to other samples.

**Fig 3.**
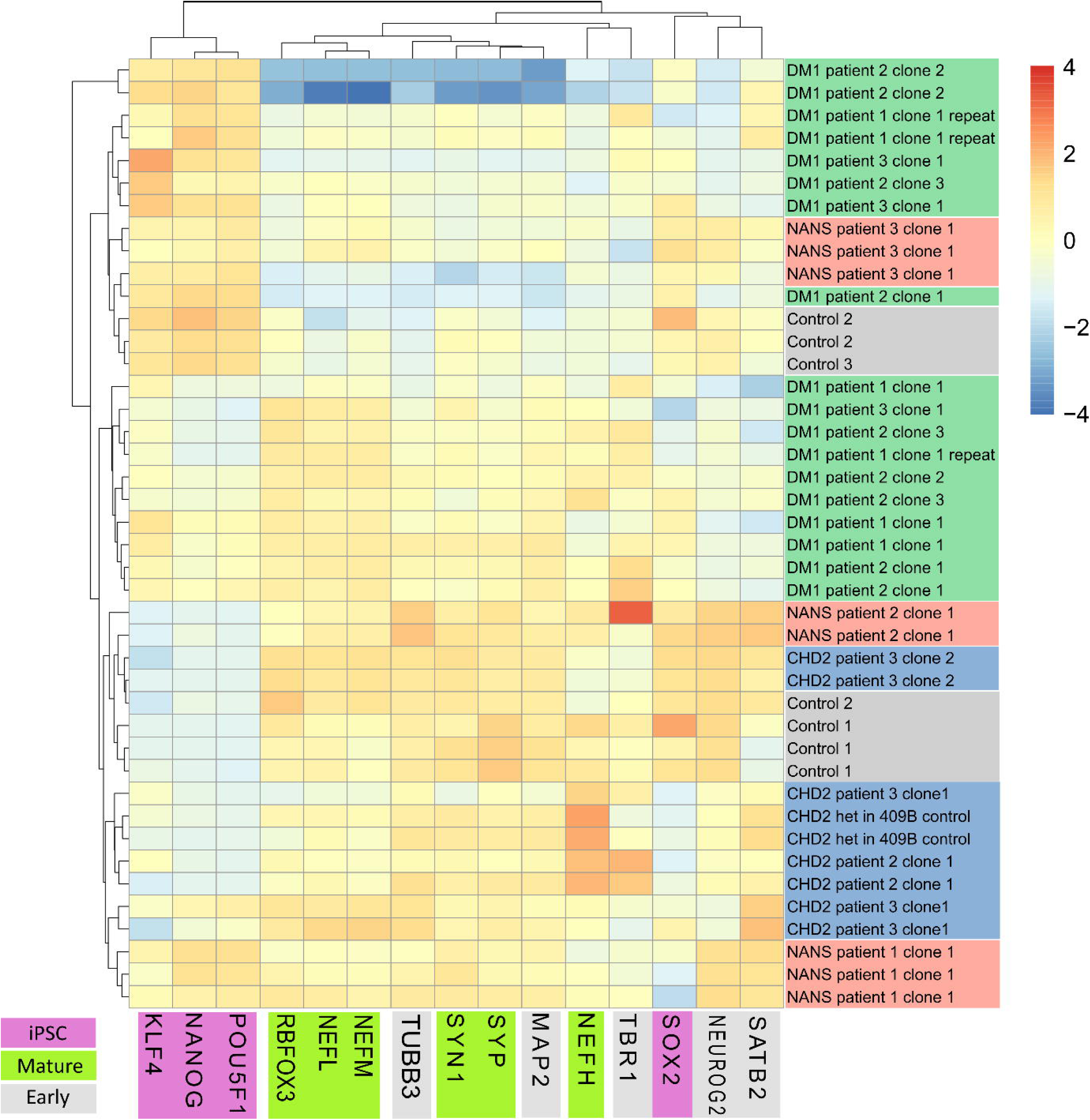
RNA levels of early and mature neuronal markers and pluripotency markers. Both the rows (samples) and columns (molecular markers) were hierarchically clustered using Euclidean distance and complete linkage clustering. Sample colors indicate the phenotype group. Marker colors indicate the type of molecular markers. The colors in the heatmap represent z-score scaled RNA expression levels of the markers.

Sample clustering based on the protein levels of these neuronal markers confirmed the finding based on transcriptomics data (Fig S9). However, the pluripotency markers (*KLF4, NANOG, SOX2 and POU5F1*) were not detectable in the proteomics measurements and thus could not be included in this analysis. We also examined correlations among neuronal marker expression levels. Since differentiation in iPSCs is driven by *Ngn2* overexpression, we were interested in whether other markers correlated with this key regulator of neuronal differentiation. As visualized by a heatmap of the correlation levels (Fig S10), neuronal markers demonstrated lower correlations with *NEUROG2* levels (human equivalent of mouse *Ngn2;* quantified levels are a mix of human and mouse) than among themselves. This suggests that other mechanisms than *NEUROG2* expression are responsible for the differences in the degree of neuronal differentiation.

In conclusion, the expression analysis of neuronal markers revealed variability among clones derived from the same donor, confirming the findings from the pairwise correlations. Moreover, the low expression levels in DM1-P2-C2 suggest that this particular iPSC line had a reduced potential for neuronal differentiation.

#### Multi-omics integration captures differences in metabolism and neuronal differentiation

To identify the largest sources of variation in the multi-omics data set, we integrated the omics layers using Multi-Omics Factor Analysis (MOFA) [20]. Using the default settings of this tool revealed a maximum set of 6 latent factors, termed as MOFA factors, that displayed shared variance across the omics layers (Fig 4A). Factor 1, represented the largest proportion of shared variation, with strong contributions from the metabolomics and lipidomics data, and modest contribution from the proteomics data. Factor 2 mainly comprised contributions from RNAseq, proteomics, EMseq and targeted lipidomics data. In contrast, factors 3–6 were largely driven by EM-seq data and explained a smaller proportion of variation, and were therefore not considered further. Since factors 1 and 2 did not separate patient and control lines (Fig. 4B), they likely captured non-disease-related variation. We therefore focused on the features with the strongest MOFA factor weights, which show how much each molecule contributes to the factors. These high-weight features helped clarify the biological meaning of the two main latent dimensions

**Fig 4.**
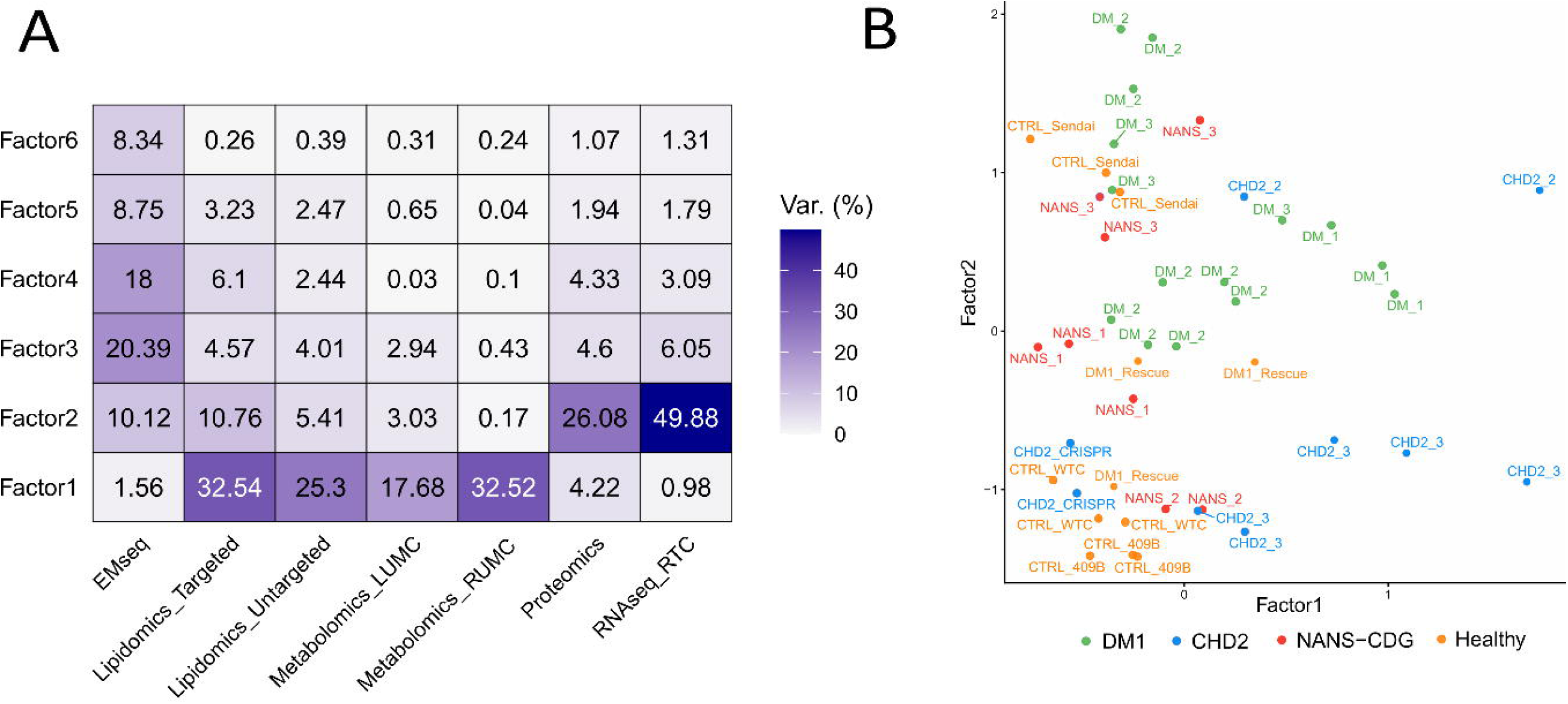
MOFA captures multi-omics variation. A) Variance contributions (in percentages) of the different omics data sets per factor. B) MOFA sample scores on factor 1 and 2, colored by disease group.

To interpret factor 1, we examined the corresponding feature weights (Fig 5(a-b), Supplementary Table 2). For the untargeted metabolomics platforms, putative manual annotation of the top features was performed by matching *m/z* values and retention times against an internal metabolite panel and the human metabolome database [21]. Several detected mass features corresponded to metabolites present in the culture medium, including three associated with glutamine and one with doxycycline. (Fig 5a). This was despite of the fact that the cell lines were cultured in the same growth media and the absence of associations between the factor scores and the sampling time or the differentiation timepoint (Fig S11). Hence, factor 1 might be indicative of the variation in how these compounds were taken up (glutamine due to active transport and doxycycline due to passive diffusion) or metabolized by the cell lines. Other features with high weights on factor 1 were identified as kinetin-9-riboside and N6-isopentenyladenosine, both of which have been implicated in cell growth pathways and mitochondrial function [22].

**Fig 5.**
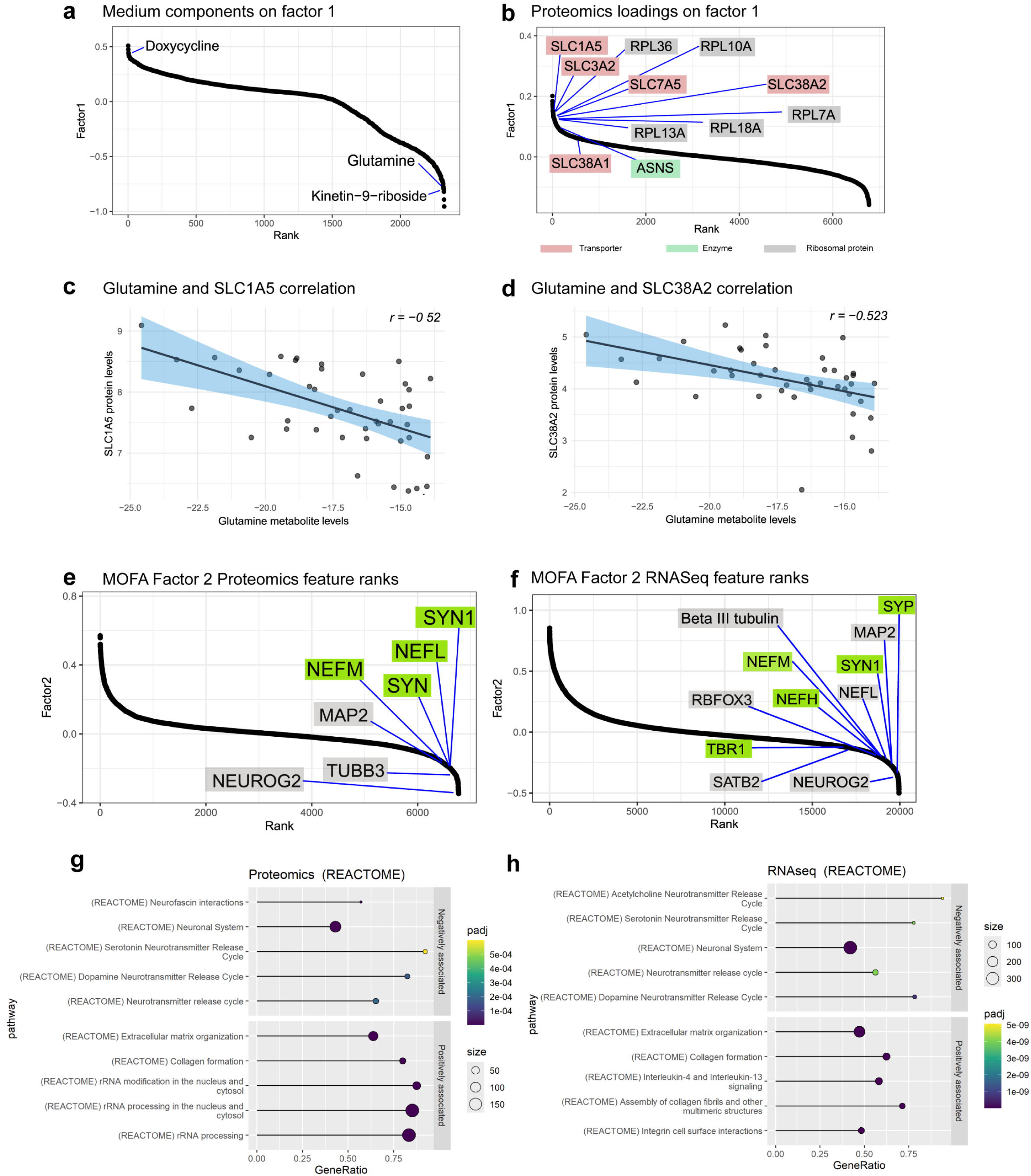
MOFA factor 1 interpretation. a) Metabolomics (LUMC) feature weights on factor 1and proteomics feature weights where some of the top loadings associated with the medium components are highlighted b) Proteomics feature weights on factor where some of the top loadings are highlighted: glutamine transporters (red), glutamine enzymes (green) and ribosomal proteins (grey) c) Spearman correlations of the measured glutamine levels and protein levels of the amino acid transporter SLCA1A5 d) Spearman correlations of the measured glutamine levels and protein levels of the amino acid transporters SLC38A2 e) RNAseq feature weights of early and late neuronal markers on factor 2. f) Proteomics feature weights of early and late neuronal markers on factor 2 g) Top 10 Reactome pathways with the highest normalized enrichment score (NES) using the proteomics data factor 2 loadings. h) Top 10 Reactome pathways with the highest normalized enrichment score (NES) using the RNASeq data factor 2 loadings.The dot size represents the size of the respective pathway and is colored according to its adjusted p-value. The GeneRatio represents the number of leading-edge genes compared to the total number of genes in the pathway.

Next, protein feature weights were used for gene set enrichment analysis (GSEA). Ranking proteins according to factor 1 revealed numerous enriched pathways across KEGG, Reactome, and WikiPathways (Supplementary Table 3). Several pathways were linked to metabolic responses to nutrient starvation, including the EIF2AK4 (GCN2) pathway activated during amino acid deficiency, which was one of the top enriched pathways. Remarkably, the activation of this pathway has been directly associated with glutamine deprivation [23,24]. In addition, the enrichment of the SLC-mediated transmembrane transport pathway can be linked to the variation in glutamine uptake, as this pathway includes known neuronal glutamine transporters, such as SLC1A5, SLC38A1 and SLC38A2 [25,26]. The negative correlation between the intracellular glutamine concentrations (Fig 5(c-d), Fig S13) and the expression of the transporter and ribosomal proteins suggest that these proteins are upregulated to compensate for low glutamine availability. Interestingly, also ASNS (asparagine synthetase), the enzyme responsible for the conversion of aspartic acid and glutamine into asparagine and glutamate, was among the top-scoring proteins on MOFA factor 1.

Other pathways associated with MOFA factor 1 were related to mitochondrial activity, such as Mitochondrial complex I assembly and Mitochondrial translation initiation. These were enriched due to strong contributions from redox-associated proteins, including catalase and several NDUF family members (e.g., NDUFV1, NDUFS1, NDUFS2). Moreover, multiple components of the mitochondrial translation machinery were enriched among the loadings, including mitoribosomal proteins (MRPL21, MRPL47, MRPS25, MRPS23, MRPS34).

Lastly, we examined the lipidomics (targeted) feature weight based on structural properties [27]. Enrichment analysis of factor 1 weights revealed strong overrepresentation of several phospholipid classes, including phosphatidylinositol (PI), phosphatidylethanolamine (PE), and phosphatidylglycerol (PG), were strongly enriched (Supplementary Table 4, Fig S14). Notably, PEs are the most abundant lipid classes on the mitochondrial membrane and have been reported to be essential for mitochondrial activity [28]. In addition, we observed significant correlations between lipid chain length and feature weights across multiple lipid classes (Fig S15-16). For instance, triglyceride (TG) chain length was negatively correlated with the MOFA feature weights (r = –0.58). Consistent with previous findings, shorter-chain TGs are indicative of higher metabolic activity, whereas longer-chain TGs are typically associated with less active cells [29]. Taken together, factor 1 represents differences in the expression of ribosomal proteins and metabolic enzymes, possibly as a consequence of glutamine starvation in certain cell lines or clones.

In factor 2, we analyzed the feature weights derived from RNA-seq, EM-seq, proteomics, and targeted lipidomics data. We first observed that the neuronal markers investigated earlier, such as *MAP2, NEUROG2, SYN1 and SYP,* (Fig 3) were strong negative contributors to factor 2, in both the proteomic (Fig 5e),and transcriptomic layers. In addition, several top negative loadings on MOFA factor 2 are proteins associated with neuronal development, including Neuromodulin, Hippocalcin, and Neogenin. Consistent with the finding from these previous analyses, the DM1-P2-C2 samples was distinct from the other two clones of the same patient, demonstrating the highest scores on MOFA factor 2 (Fig 4B). We next performed GSEA using RNA-seq, proteomics, and EM-seq feature weights. For the latter, we linked all CpG sites with the nearest gene, to map these methylation sites to biological pathways. GSEA revealed numerous significantly enriched neurological pathways across these datasets (Fig 5(g-h), Supplementary Table 5).

Finally, we examined the lipidomics feature loadings using the same overrepresentation analysis. This revealed strong positive enrichment of triglycerides, sphingolipids (SM), and hexosylceramides (HexCer) on factor 2 weights (Supplementary Table 4, Fig S14). Notably, previous studies [30] also reported high abundance of these lipid classes in neurons, supporting our observations. Taken together, these results suggest that MOFA factor 2 captures variation in the neuronal differentiation stage among the different cell lines.

In conclusion, integration with MOFA revealed the major sources of variation within our multi-omics dataset generated from patient lines from multiple disease groups. These variations were not directly linked to the specific disease phenotypes but instead reflected other biological differences among the iPSC lines, including metabolic activity and neuronal differentiation. Consequently, the scores and feature weights of these factors may serve as useful covariates to correct for such differences, thereby promoting the discovery of robust and disease-specific signatures.

### Identifying disease-specific signatures in the multi-omics data

#### Linear models identify differential omics features for each patient group

Our next aim was to identify disease-specific signatures within each patient group that are robust against the observed variations. To determine the most differential omics measurements for the different patient groups, we applied linear mixed models. These models account for the sample hierarchy in our dataset, control for intra-donor variation and make sure that clones and biological replicates are not seen as completely independent observations. Specifically, we included the sample and clone numbers as random effects, where clones are nested within samples (see Methods).

We ran these models for each individual omics feature, comparing patient groups against control lines. For the DM1 patient lines, the CRISPR Rescue line (Fig S1) was included as a control. Additionally, we performed a separate analysis limited to DM1 patients 2 and 3, since patient 1 was reported with a much milder phenotype due to a shorter disease-causing CTG repeat. We also conducted models comparing all patient lines combined with controls, as well as models testing for sex-associated effects. The number of significant associations after adjusting for multiple testing are summarized in Figure 6 and Supplementary Tables 6-11. Overall, the proportion of significant associations was maximally 1.5% of features per dataset. This is expected, as the small number of patients per group reduces statistical power. Consequently, the p-values are more conservative than those obtained from fixed-effects models (Fig S17) that falsely treat all samples as independent.

**Fig 6.**
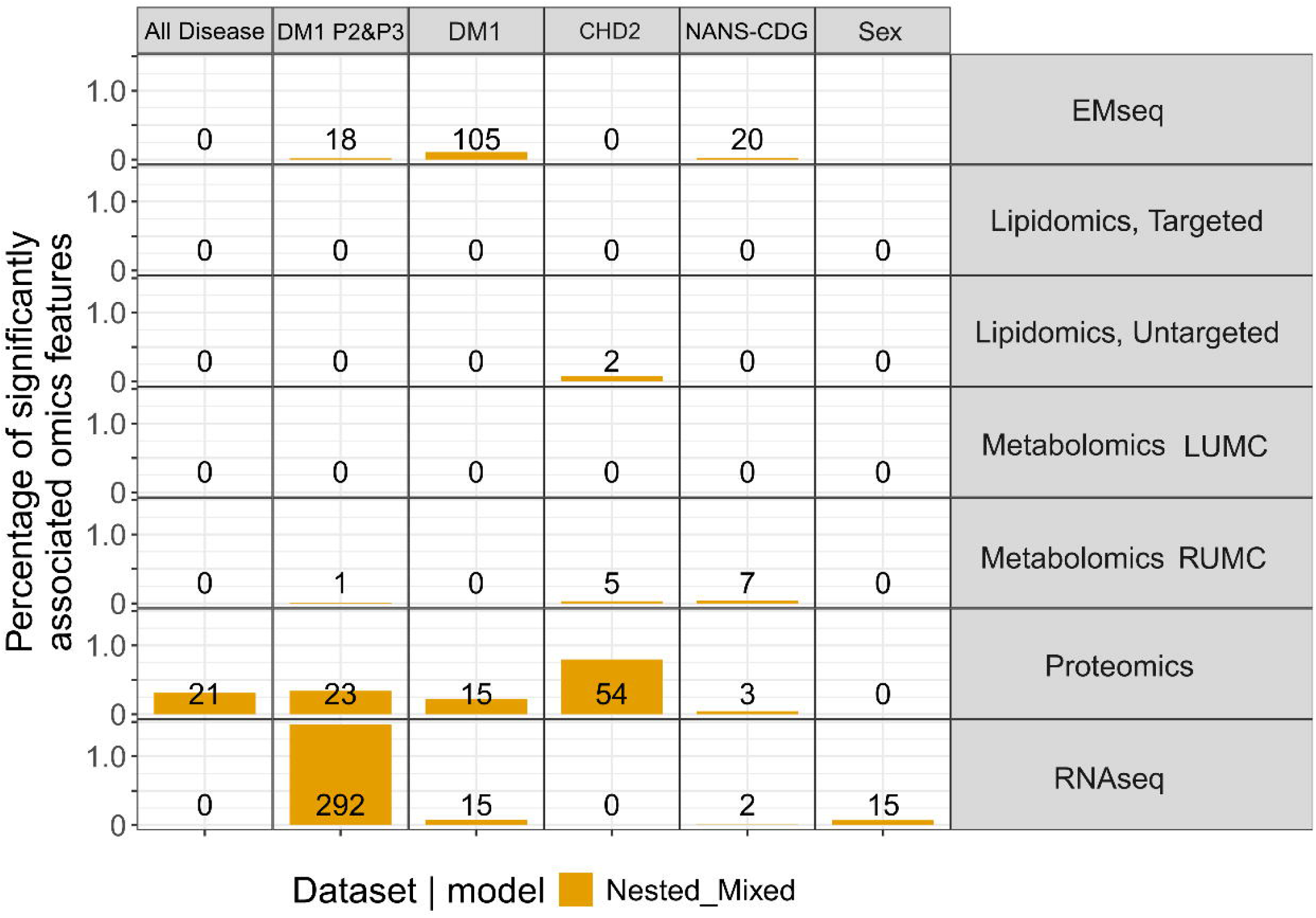
Significant associations between omics layers and disease groups found with linear mixed models. The yellow boxes represent the percentage of significant associations per omics data set (right), per contrast (top, all disease lines *vs.* controls, DM1 P2 and P3 *vs.* controls, all DM1 patients *vs.* controls, CHD2 patients *vs.* controls, NANS-CDG patients *vs.* controls and males *vs.* females). The absolute numbers of significant associations are presented inside the yellow boxes.

Proteomics data consistently exhibited the strongest and most significant associations with disease groups. As expected due to the more severe phenotype, a substantially larger number of RNA associations were identified when focusing only on DM1 patients 2 and 3 instead of the whole group of DM1 patients. When comparing all disease lines together with the control lines, only a few proteomics specific associations remained, indicating limited similarity for these disease groups at the molecular level. Finally, also few associations with sex were found, suggesting that the sex of the donor and the presence of an additional X or Y-chromosome has a small influence on the omics profiles of the iPSC models.

Some known disease signatures were rediscovered by the nested mixed-effect model. For instance, in the RUMC metabolomics dataset, the fourth-ranked metabolite corresponded to a significant upregulation of N-acetylmannosamine-6-phosphate (ManNAc-6-P) in the NANS-CDG patient lines (Fig S18), a previously described biomarker of this disorder [18]. Moreover, several proteins (e.g., mismatch repair (MCM) family genes) that were significantly associated with the *CHD2* patient lines, have a function in DNA repair mechanisms, a link that has been reported before [31]. Hypermethylation of the CTG-repeat containing region of the *DMPK* gene, of which its expansion is causing DM1 and in which hypermethylation was observed in patients with long repeats [32], was observed in the DM1 cells from patients 2 and 3 with the longer repeats, but not in cells from patient 1 with the shorter repeat expansion (Rahm *et al.*, manuscript in preparation).

#### Correction for MOFA factor 1 and 2

Ultimately, we aimed to minimize the effect of cell line variability on the identification of possible disease markers by correcting for MOFA factor 1 (metabolic/mitochondrial variability) and factor 2 (neuronal differentiation variability). To do this, we regressed out the MOFA samples scores on factor 1 and 2, and ran the same nested linear mixed models with the residuals resulting from the regressions (Fig S19, Supplementary Tables 12-17). This resulted in a larger number of disease-associated metabolites, proteins, and transcripts for CHD2 and NANS-CDG patient lines. In contrast, DM1-associations were limited after correction for the MOFA factors. This could partially be explained by the large degree of variation among the DM1 clones on these factors, which may have resulted in false positives without correction. Again, some of the known disease-specific associations remained after correcting for the MOFA factors, for example, ManNAc-6-P for the NANS-CDG patient lines. This reinforces the confidence in the workflow presented in this study and suggests that other differentially expressed omics features may represent candidate biomarkers for the individual disease groups.

We inspected the omics features that were only significant after correction for the MOFA factors 1 and 2. As expected, the number of significant associations increased, because standard deviations decreased after correction, in particular in the groups of healthy control lines (Fig. S20). Among the features that were differentially expressed in the CHD2 after correction were the DNA replication licensing factor MCM4 and the ribosomal protein RPL35, linked to the affected DNA repair and ribosomal translation pathways in CHD2-deficient cells, respectively. Interestingly, L-aspartic acid, which can act as a neurotransmitter, was more significant and showed a larger difference between CHD2-deficient lines and controls after correction. Together, these findings indicate that the correction improved the detection of disease-specific variation.

### Identifying possible causes for interclonal differences

#### Differentially methylated regions

We explored (epi)genetic factors that may contribute to the observed differences in the DM1-P2 clones. Previous studies have suggested that incomplete erasing of DNA methylation during iPSC generation may contribute to differences in DNA methylation patterns in iPSC lines [13,33,34]. To explore this, we analyzed differentially methylated regions (DMRs) across clones. Using the dispersion shrinkage method for sequencing data (DSS) [35], we identified 2,428 DMRs when comparing DM1-P2-C2 with the other clones from the same patient (Supplementary Table 18). For reference, the corresponding analyses for clone 1 and clone 3 yielded 915 and 2,115 DMRs, respectively. Strikingly, many of the DMRs in clone 2 clustered within a single locus on chromosome 12, spanning several protein-coding genes (*TMEM132D*, *TMEM132C, RIMBP2, GLT1D1*, and *TMEM132B*). Out of these, 168 DMRs mapped to genes of the Transmembrane Protein 132 (*TMEM132*) family. This gene family has been implicated in several neurological disorders, including panic disorder, anxiety disorder, and depression [36,37]

Closer inspection of the largest DMR in the *TMEM132D* gene demonstrated that this clone clustered with other lines hypo-methylated in this region (Fig S21), whereas clones 1 and 3 from this patient clustered with other DM1 patients and other hypermethylated lines. Methylation of this region is therefore highly variable between clones, irrespective of the donor where they are derived from. Interestingly, these differences were reflected on the transcriptomic level. For instance, RNA levels of *TMEM132D* were higher in clone 2 than in clones 1 and 3 (Fig S22). More generally, the samples exhibiting the highest methylation levels in this region showed lowest expression of *TMEM132D.* The expression levels of another TMEM132 family member, *TMEM132E*, were reduced in DM1-P2-C2. This corresponds to the MOFA results, where the *TMEM132E* transcript was one of the top contributing loading on MOFA factor 2. In conclusion, these findings indicate a direct link between seemingly stochastic DNA methylation and transcriptional regulation in these loci, possibly affecting neuronal differentiation.

#### Genetic differences at the iPSC stage

To evaluate whether stochastic differences in methylation present prior to differentiation could contribute to the variation observed among the DM1-P2 clones, we generated additional EMseq data sets from the different clones of DM1-P2 in undifferentiated iPSCs. Already in the undifferentiated state, pronounced DNA methylation pattern differences were present between the clones (Fig 7). The differences between clones were observed to be larger than the differences between differentiated and undifferentiated cells derived from the same clone. This suggests that 7-day differentiation has a lower impact on DNA methylation than the reprogramming to iPSCs. In addition, the correlation between clone 2 iPSC and its corresponding iNeuron was higher (∼0.9) than for the other two clones (∼0.8). This is consistent with the lower degree of differentiation of clone 2 after 7 days of differentiation compared to clones 1 and 3.

**Fig 7.**
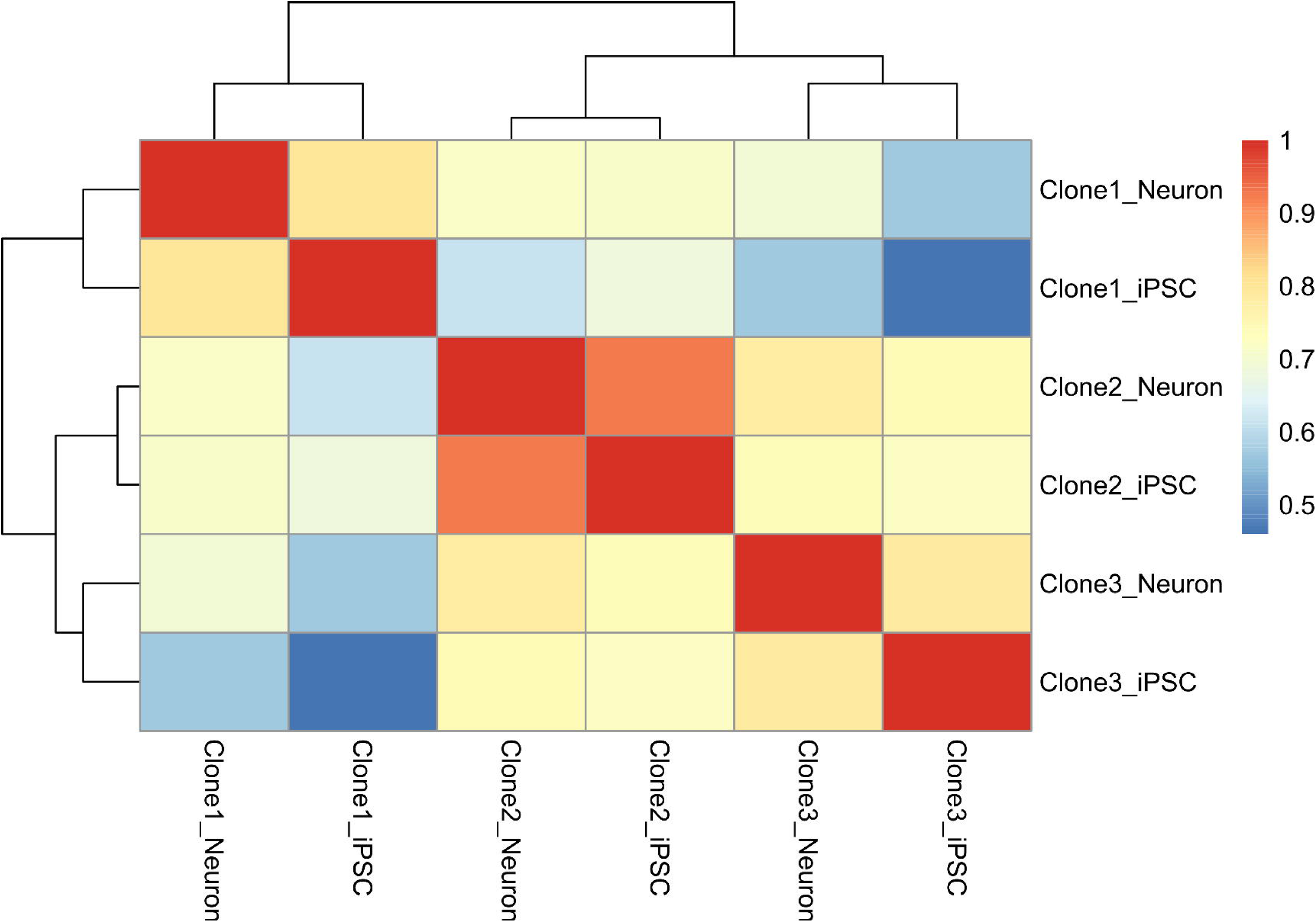
Pairwise correlations of the DM1 P2 EM-seq data sets measured at the iPSC stage and the iNeuron stage. The heatmap represents Spearman’s correlations (see legend) of the set of 100,000 most variable CpG sites that were used throughout this study.

## Discussion

Our study provides insight into the variability inherent to iPSC-based disease models. With our experimental design, we were able to pinpoint the major contributors to this variability as reflected in the multi-omics data. Neuronal differentiation status was an important source of variation, even for clones of the same donor. Moreover, variation in nutrient uptake and metabolism had a large impact on the omics profiles and may cause differences in metabolic activity between iPSC-derived neuronal cell cultures. Despite the observed variability, we were able to correct for the major sources of variability and recover known and novel disease-associated molecular signatures for all three patient groups.

Our study has important implications for the design of future studies in iPSC-derived neurons. We observed high variability between patients in the same disease group, sometimes even comparable to the differences between disease groups. Notably, the variability between clones from the same patients was comparable to the variability between patients. On the contrary, biological replicates of the same cell culture showed high consistency in all omics measurements and were in some omics layers comparable to correlations between technical replicates. This is likely due to the fact that all the cultures were processed by the same experimenters in the same laboratory. Differences between experimenters and laboratories may introduce additional variation in the omics profiles. However, in the context of an experiment in a single laboratory, resource- and budget-efficient experimental designs aimed at the discovery of disease-specific variation should preferentially include more different patients or more different clones from the same patient over biological replicates of the same clones.

The high disease-unrelated variability in iPSC-derived models confirms observations in previous studies [6,7,9,10,12,38,39]. These previous studies attributed most of this variation to genetic differences that are already present at the iPSC stage. However, our results demonstrate that clones derived from the same patient display variability comparable to that observed between different patients, particularly at the transcriptomic and proteomic levels, contradicting previous studies [40]. This indicates that other factors during reprogramming, clonal selection, or differentiation can contribute to cell line differences. The low correlation of EMseq data measured among different clones from the same patient at the iPSC stage confirmed these differences. Epigenetic differences, in turn, could partially explain the clonal differences found in the other omics data sets.

Further, DM1-P2-C2 showed reduced levels of neuronal markers compared to the other clones from this patient, and higher levels of pluripotency markers. This illustrates how different the neuronal differentiation speed or efficiency can occur among clones of the same donor. This difference cannot be explained by variances in lab conditions, because all clones were cultured and processed in parallel. Moreover, cell density was similar in all culture dishes. We therefore speculate that these differences may, at least partially, be explained by differences in DNA methylation. We observed a large number of DMRs between clones from the same patient that were also linked to changes in the gene expression levels of the corresponding genes. The clonal differences in DNA methylation were already present at the iPSC stage and may be the consequences of stochastic differences in the erasing and reestablishment of DNA methylation patterns during reprogramming [41,42]. Finally, the differences in DM1-P2-C2 could also be caused, in part, by genomic aberrations that arose during cell line generation, or were already present in the original tissue as mosaicisms, which is not unusual in fibroblast [43].

Multi-omics integration using MOFA identified latent dimensions across the metabolomics, lipidomics and proteomics data (Factor 1), and across RNAseq, proteomics, EMseq and lipidomics (Factor 2) that explain a large proportion of the total variation in all these data. These multi-omics factors were useful in explaining the variability within iPSC models, and revealing insights that single-omics analyses alone could not capture. More importantly, we have shown that information from one omics layer can be used to correct or refine another, highlighting the unique advantages of multi-omics studies.

Factor 1 reflected large differences in intracellular glutamine concentrations, which may result from altered expression or activity of glutamine transporters or metabolic enzymes that we found in the proteomics data. Reduced intracellular glutamine is known to activate the EIF2AK4 (GCN2) amino-acid starvation pathway, leading to phosphorylation of eIF2α and suppression of protein translation [24–26]. In addition, factor 1 also showed variation in intracellular doxycycline concentrations, which can specifically inhibit mitochondrial protein translation [44]. Thus, the observed upregulation of amino-acid transporters and ribosomal protein genes may represent a compensatory response for these stress conditions. Altered glutamine-to-glutamate ratios could affect glutamatergic neuronal metabolism. However, the impact on glutamatergic signaling and neuronal function would need further studies.

MOFA Factor 2 was strongly associated with neuronal differentiation status, with neuronal markers highlighted as top contributors. Additional high-loading features on this factor may also represent novel indicators of the differentiation process. For example, opposite to the neuronal markers, Neuroblast differentiation-associated protein (AHNAK) was among the top positive contributors to this factor, suggesting that this protein is more abundant in less differentiated neurons. We therefore propose further investigation of this set of potential markers, which could aid researchers in prioritizing iPSC clones with higher differentiation potential. Moreover, the expression of omics features that score high on factor 1 and 2, can be used to correct for differences in metabolic activity and neuronal differentiation differences, when large-scale multi-omics data are not available. Instead of residualizing the scores on the MOFA factors, it would be possible to perform regression of smaller subsets of biomarkers to control for unwanted variability, although this is likely less robust than regression of the MOFA factors.

Using nested linear mixed models, we identified the most discriminative features for each patient group. Some of these biomarkers were previously reported by other studies. For example, accumulation of ManNAc-6-phosphate levels was already associated with NANS-CDG [18], which was also found in our metabolomics data as fourth-ranked metabolite associated with the NANS cell lines. Additionally, our study identified more than 100 highly differential omics features in different omics layers that were not associated with *NANS-* and *CHD2-*related disorders before and thus hold potential as biomarkers for disease and justify further investigation.

This study has certain limitations that should be considered. Notably, we analyzed only three patients per disease group. Ideally, a larger sample size would allow for more accurate conclusions on the variability between patients, clones, and replicates. However, as DM1, CHD2, and NANS-CDG are rare diseases, the sample size is inherently limited. This emphasizes the importance of studying the reproducibility of iPSC studies and the need for correction of disease-unrelated variation.

Furthermore, the cell cultures were not all created from the same tissue types. The CHD2 patient models were generated based on PBMCs, whereas for the other lines, fibroblasts were used to reprogram to iPSCs. This can specifically affect RNA and DNA methylation levels, as shown before [45,46]. Therefore, differences between CHD2 patients and controls or other disease lines may possibly be confounded by the original cell types. We have mitigated this by including one disease line where a *CHD2* mutation is introduced into a control line, but results should be interpreted with care.

Finally, correcting for MOFA the factors may unintendedly remove biologically relevant information. Although these factors did not separate patient groups and represent orthogonal, uncorrelated dimensions, we cannot fully exclude the possibility that such corrections also removed disease-related biological signals.

Taken together, this study provides guidelines for future multi-omics studies based on iPSCs and specifically iPSC-derived neurons:

- We stress the need for lines from multiple patients, and multiple clones per patient, whereas measurements of independent cultures from the same clone are less important.
- Sample concordance checks should be standard procedures of omics data studies, especially in studies with complex experimental design like in this study.
- Case-control comparisons in studies with iPSC-derived neurons can be confounded by differences in amino acid starvation, protein translation and mitochondrial activity and may require methods to correct for these differences.
- We suggest to prioritize proteomics over other omics analyses in studies investigating neurodevelopmental disorders, given the higher numbers of significant disease-relevant features and higher interpretability of the results.

## Methods

### Human iPSC lines and CRISPR/Cas9 edited iPSC lines

#### Control lines

One control line (WTC) in this study, USFi001-A (sourced from the Coriell Institute, GM25256, RRID: CVCL_Y803), was reprogrammed with non-integrating episomal vectors from skin fibroblasts of a healthy 30-year-old male. The second control line (409B) used is BRCi023-A (sourced from the Center for iPS Cell Research and Application, Kyoto University; Kyoto; Japan; RRID: CVCL_K092). It was reprogrammed with non-integrating episomal vectors from skin fibroblasts of a healthy 36-year-old female. The third control line (Sendai) control, UMGi020-A sources from the University Medical Center Goettingen (CVCL_A4KA), was obtained from a 24-year-old female subject.

#### CHD2-lines

Peripheral mononuclear blood cells (PBMCs) were isolated from patients via blood samples taken during routine diagnostic testing. One patient line was generated using CRISPR/Cas9 gene editing. A heterozygous indel variant in exon 4 of CHD2 was introduced in an iPSC line derived from control line 409B. A short guide RNA (sgRNA) was designed (ATCCTGATGTTTATGGGGTC) and cloned into pSpCas9(BB)-2A-Puro (PX459) V2.0 (Addgene, #62988) according to previous studies. 8×10^5^ single iPSCs were nucleofected with 5 µg of the generated pSpCas9-sgRNA plasmid using the P3 Primary Cell 4D-Nucleofector Kit (Lonza, #V4XP-3024) in combination with the 4D Nucleofector Unit X (Lonza, #AAF-1002X). After nucleofection, cells were resuspended in E8 Flex supplemented with RevitaCell (Thermo Fisher Scientific, #A2644501) and seeded on biolaminin 521 (Biolamina, #LN521) pre-coated wells. 24 hours after nucleofection, 0.5 µg/ml puromycin was added for 24h. Puromycin-resistant colonies were manually picked and sent for Sanger Sequencing to ensure heterozygous editing of exon 4. Genomic stability was assessed by detection of recurrent abnormalities using the iCS-digitalTM PSC test, provided as a service by Stem Genomics (https://www.stemgenomics.com/).

CHD2-deficient iPSC lines were cultured on biolaminin 521 (Biolamina, #LN521), and CtrlAn/An lines under feeder-free conditions either on Matrigel (Corning, #354277) coated plates or 0,125mg/cm2 of iMatrix-511 (ReproCell, #NP892-012). Cells were passaged before reaching 70% confluency using an enzyme-free reagent (ReLeSR, Stemcell, #05872) or a 0.5 mM EDTA solution and not kept for more than 10 passages. All iPSCs were regularly tested for mycoplasma contamination using MycoAlert PLUS (Lonza, #LT07-703) and their identity was confirmed by short tandem repeat profiling.

#### NANS-lines

Three NANS-CDG patient iPSC lines were generated from skin fibroblasts. NANS-CDG patient 1 (Radboudumc, iPSC nr: IPS17-00058), NANS-CDG patient 2 (Radboudumc, iPSC nr: IPS17-00057) and NANS-CDG patient 3 (Radboudumc, iPSC nr: IPS17-00045) were used in this study. All iPSCs were cultured on Matrigel (Corning cat. No. 356237) or human recombinant laminin 521 (BioLamina, cat no. LN521-02) in Essential 8 flex medium (Thermo Fisher Scientific, cat. No. A2858501) supplemented with primocin (0.1 μg/mL, Invivogen, cat. No. ant-pm-2), low puromycin (0.5 μg/mL, Sigma-Aldrich, cat. No. P9620), and G418 (50 μg/mL, Sigma-Aldrich, cat. No. G8168) at 37°C and 5% CO2. Medium was refreshed every 2 to 3 days and cells were passaged weekly, depending on confluency, using an enzyme-free reagent (ReLeSR, Stem Cell Technologies cat. No. 05873), with a maximum passage number of 20. Cells were frequently tested for mycoplasma infection. The cell line was not authenticated.

#### DM1-lines

All DM1 lines were generated from skin fibroblasts by lentiviral transduction at the Radboudumc Technology Center Stem Cells (Nijmegen, The Netherlands). One of the patient lines was used for CRISPR/Cas9 editing to generate an isogenic control. The expanded CTG repeat was excised by using two guide RNAs (5’-GCTCGAAGGGTCCTTGTAGC-3’) (5’-GCTGAGGCCCTGACGTGGAT-3’) as previously described [16,47]. The double-strand break was repaired by non-homologous end-joining, yielding a clone that showed an excised allele without indels and an unedited healthy allele (CTG5). Off-target analysis was performed by sequencing the top five off-target sites for each guide RNA that were predicted by CRISPOR [48] and Benchling (Biology Software). Digital droplet PCR of the most common 24 copy number variations did not reveal any variations in this CRISPR-edited clone.

### Generation of rtTA-Ngn2-positive iPSCs and their differentiation towards glutamatergic neurons

To differentiate iPSCs into glutamatergic neurons, an inducible Ngn2 gene cassette was introduced into iPSCs following a previously established protocol [49]. The iPSCs were transduced with lentiviral vectors to stably integrate the rtTA (pLV-EF1α>Tet3G:IRES:Neo) and Ngn2 {pLV[TetOn]-Puro-TRE3G > mNeurog2([NM_009718.3])} transgenes into their genome. Forty-eight hours post-transduction, cells carrying both vectors were selected using G418 (25 μg/ml; Sigma-Aldrich, #G8168) and puromycin (0.5 μg/ml; Sigma-Aldrich, #P9620), with the concentrations gradually increased over time. The iPSCs that survived the selection were cultured in Essential 8™ Flex Basal medium (Gibco) supplemented with primocin (Invivogen, #ant-pm-2), G418 (50 μg/ml), and puromycin (0.5 μg/ml) on Geltrex-(Gibco)-coated plates at 37°C/5% CO2. Cells were passaged once or twice a week with either ReLeSR (STEMCELL Technologies) or UltraPure^TM^ 0.5M EDTA, pH 8.0 (final concentration 0.5mM; Invitrogen) when they reached 80%–90% confluence. The iPSCs were then differentiated, according to protocol, for 7 days into cortical glutamatergic neurons through doxycycline-inducible *Ngn2* overexpression. Primary rodent astrocytes were not added to the cultures.

### Sample collection protocols for the different omics data sets

#### EMseq

Medium was aspirated and cells were washed once with 1XDPBS at RT. Cells were scraped in 300 μl 1XDPBS and transferred into a 1.5 ml Eppendorf tube. Eppendorf tubes were centrifuged (1000RPM, 5 min), supernatant was aspirated, and cells were snap frozen in liquid nitrogen. Cell pellets were stored at −80°C until further processing. DNA isolation was performed with the QIAmp® DNA Mini Kit (Cat. #51304 and 51306), following the standard protocol.

#### Transcriptomics

Medium was aspirated and cells were washed once with 1XDPBS (Gibco; DPBS, no calcium, no magnesium; Cat. #14190144) at room temperature (RT). 300 μl of 1X DNA/RNA Shield Stabilization Solution (ZYMO Research; Cat. #R1100-250) was added and cells were scraped with a cell scraper. Cell lysates were transferred into a 1.5 ml Eppendorf tube, snap frozen in liquid nitrogen and stored at −80°C until further processing. RNA isolation was performed with the Quick-RNA Microprep Kit (ZYMO Research; Cat. #R1050), following the standard protocol.

##### Proteomics

Medium was aspirated and cells were washed once with 1XDPBS at RT. Cells were scraped in 300 μl 1XDPBS and transferred into a 1.5 ml Eppendorf tube. The well was rinsed with 1 ml 1XDPBS. Eppendorf tubes were centrifuged (max speed, 5 min), supernatant was aspirated, and cells were snap frozen in liquid nitrogen. Cell pellets were stored at −80°C until further processing.

##### Metabolomics

Medium was aspirated and cells were washed once with ice cold 1XDPBS. Cells were scraped in 300 μl cold 1XDPBS, transferred into a 1.5 ml Eppendorf tube and centrifuged (16,100g/max speed, 30 seconds). Supernatant was aspirated, cell pellets were snap frozen in liquid nitrogen and stored at −80°C until further processing.

##### Lipidomics

Medium was aspirated and cells were washed once with ice cold 1XDPBS. Cells were scraped in 300 μl cold 1XDPBS. Cells were transferred into a 1.5 ml Eppendorf tube and centrifuged (16,100g/max speed, 30 seconds). Supernatant was aspirated, cell pellets were snap frozen in liquid nitrogen and stored at −80°C until further processing.

### Omics data measurements

#### CNV-WES

Remaining iPSCs that were not used for neuronal plating were collected into a 1.5-mL Eppendorf tube and centrifuged (1000 rpm, 5 min). The supernatant was aspirated, and the resulting pellet was snap-frozen in liquid nitrogen and stored at −80 °C. DNA was isolated using the QIAmp® DNA Mini Kit and 2.5 μg DNA. Each of the 16 iPSC cell lines were sent for WES analysis before differentiation. Sequencing was performed using the NovaSeq6000 platform. Exome enrichment was achieved using Twist-Exome 2.0 and Comprehensive Exome Spike-in Kit. Read alignment was performed with Burrows-Wheeler Aligner [50] using reference genome GRCh37/hg19. The resulting data was analyzed for CNVs using Conifer [51] and ExomeDepth (ED) pipelines [52].

#### EMseq

Sequencing libraries were generated using the NEBNext Enzymatic Methyl-seq sample preparation kit (cat# E7120S/E7120L, NEB) with unique dual indexes. Input for library preparation was 100 ng for all samples except for: CHD2-p3c2-1 (200ng), DM1-p1c1-2 (65ng), CHD2-p2c1-1 (87ng). The library preparation was performed according to the manufacturers’ instructions (manualE7120), including the addition of control DNAs (Puc19 and Lambda).

Sequencing was performed on an Illumina NovaSeq6000 system, with a paired-end 150 bp read configuration (Illumina, San Diego, CA, USA). All libraries were sequenced to a minimum of 300 M paired-end reads per sample with read length 2×150 bp. Sequencing data was demultiplexed and converted from base calls into FASTQ files using Illumina bcl2fastq2 Conversion Software v2.20.0.422. Quality control after sequencing was performed using the in-house quality control tools CheckQC (https://github.com/Molmed/checkQC) and Seqreports (https://github.com/Molmed/seqreports) as well as FastQ Screen from Babraham Bioinformatics (https://www.bioinformatics.babraham.ac.uk/projects/fastq_screen/).

Primary methylation sequencing data analysis was performed with nf-core analysis pipeline methylseq version 1.6.1 (https://github.com/nf-core/methylseq), the reference genome used was Ensembl GRCh38, release 105. Pipeline output includes alignment to the reference genome, methylation calls and quality metrics. The controls pUC19 and lambda were aligned using Bismark version 0.23.0 (Babraham Bioinformatics) and the methylation rate was compared against the threshold provided by the vendor.

We used the methrix v 1.18.0 R/Bioconductor package for data processing using BSgenome. Hsapiens.NCBI.GRCh38 as the reference genome. Methylation sites with a read coverage below 5 were set to NA. The resulting object was filtered by removing all CpG sites covered in <20% of the samples. The sites overlapping with common SNPs as defined by a minor allele frequency > 0.01 in the 1kGenomes based on MafDb.1Kgenomes.phase3.hs37d5 data package were removed using the *remove_snps* function as implemented in the methrix R package, leaving a total of 23,392,898 CpG sites. To avoid this large number of CpG sites dominating the multi-omics analysis, we selected the 100,000 most variable CpG sites, based on standard deviation.

#### Transcriptomics

RNA sequencing was performed at the Utrecht Sequencing Facility (Utrecht, The Netherlands). 100ng of total RNA was used to prepare Illumina TruSeq Stranded mRNA libraries following the manufacturers protocol, half the reagents volume, with custom 384 xGen UDI-UMI adapters from IDT. Sample libraries were pooled equimolar. Libraries were sequenced on a Nextseq2000 (Illumina) by using a P3 flowcell with 50bp paired-end reads.

Primary sequencing data analysis is performed using the RNA-seq pipeline of the Radboud Technology Center for Bioinformatics (gitlab.cmbi.umcn.nl/rtc-bioinformatics/rna-dxp-dsl2). This pipeline was used to process fastq files with alignments to the reference genome (GRCh38) and various quality metrices were analyzed. This pipeline was run using Nextflow version 22.04.5. Consequently, reads counts were filtered for a minimum coverage of x per y samples, and the resulting read counts were normalized using DESeq’s (version 1.44.0.) variance stabilization.

#### Proteomics

Proteomics measurements were performed at the Radboud Technology Center for Proteomics. Cell pellets were resuspended in lysis buffer 1:1 consisting of 8M urea 10mM Tris-HCl (pH 8) prior to reduction and alkylation of cysteine residues by addition of 1µl 50mM dithiothreitol and 1µl 50mM chloroacetamide and subsequent incubation at room temperature for 30 minutes in the dark, respectively. Next, samples were diluted 1:3 with 50mM ammoniumbicarbonate (pH 8) and 1 µl Trypsin (0.2µg/µl) was added for overnight digestion at 37°C. For each sample, 200ng peptides were loaded onto Evotip Pure tips (EV2013, EvoSep) according to manufacturer’s recommendations. Tryptic digests were analyzed by nanoflow liquid chromatography (Evosep One, Evosep Biosystems) coupled online to a trapped ion mobility spectrometry - quadrupole time-of-flight mass spectrometer (timsTOF Pro2, Bruker Daltonics) via a nanoflow electrospray ionization source (CaptiveSprayer, Bruker Daltonics). Tryptic peptides were separated by C18 reversed phase liquid chromatography (Evosep EV1137 30SPD performance column; 150 mm length x 0.150 mm internal diameter, 1.5µm C18AQ particles) using the pre-programmed 30 samples per day (30SPD) Evosep One method. The mass spectrometer was operated in positive ionization mode using the default data independent acquisition - Parallel Accumulation SErial Fragmentation (dia-PASEF) instrument method: 0.6 - 1.6 1/K0 mobility range, 100 - 1700 m/z mass range, 100ms accumulation time, 100ms ramp time, 26Da mass width, 1 Da mass overlap, 32 mass steps per cycle, 0 mobility overlap, 1 mobility window. Acquired spectra were streamed directly to ProteoScape (v2025b, Bruker Daltonics) for protein identification against the Uniprot human protein sequence database (downloaded Jan 2024) and subsequent label-free quantitation using the following settings: Spectronaut v19 directDIA+ (Fast) workflow, 0.2 precursor PEP cutoff, 0.01 precursor Q-value cutoff, 0.01 protein Q-value cutoff global, 0.01 protein Q-value cutoff, 0.75 protein PEP cutoff, full tryptic specificity, allowed up to 2 missed cleavages, carbamidomethyl (C) as fixed modification and Oxidation (M) as variable modifications, protein group specific peptides were used for quantitation.

Features with missing values in more than 20% of the samples were removed. The remaining missing values were replaced by half of the minimum value of the respective column. Measurements of technical replicates were merged by taking the mean. Log2-transformation was applied to the resulting intensity values.

#### Metabolomics (RUMC)

Untargeted metabolomics measurements at the Radboud university medical center (RUMC) were performed using ultra-high-performance liquid chromatography mass spectrometry (UHPLC-MS). To extract metabolites, cells (neurons at DIV 7) were quickly washed twice with room-temperature washing buffer, consisting of ammonium carbonate (75 mM, Honeywell, cat. No. 207861-500G) dissolved in MilliQ, with the pH set to 7.2–7.4 using acetic acid. Immediately after removal of the washing buffer, the cells were snap-frozen in liquid nitrogen. Next, metabolites from frozen cells were extracted with 700 μL cold (−20°C) 2:2:1 (v/v/v) methanol:acetonitrile:water (Honeywell, cat. No. 34860–2.5 L/Biosolve, cat. No. 0001207802BS) for 4 min. The supernatant was collected while it was kept cold and the extraction was repeated with another 700 μL cold extraction solvent for 4 min. The two extracts were pooled and centrifuged at 14 000 x g for 3 min, at 4°C. Supernatants were dried using a vacuum centrifuge (SpeedVac, Thermo Fisher Scientific). Metabolites were reconstituted in 100 μL MilliQ (with 0.1% fortmic acid) and transferred to 96-well plates for LC-MS analysis, or stored at −80°C until use. Untargeted metabolomics measurements were performed using an Agilent 1290 UHPLC system coupled to an Agilent 6545 QTOF mass spectrometer, equipped with a dual electrospray ionization source. Each sample was run in duplicate in both negative and positive ionization modes. Two microliters of extracted metabolite sample were injected onto an Acquity HSS T3 (C18, 2.1 × 100 mm, 1.8 μm) column (Waters) operating at 40°C. Each analytical batch included neuronal samples, analytical quality control samples, and a pooled sample of all neuronal samples to check for integrity of the computational data preprocessing pipeline. To correct for possible run-order influence on signal intensities, the technical duplicates were analyzed in antiparallel run order. The acquired raw LC/MS data were processed using our in-house computational pipeline to perform peak detection and retention time alignment [53], generating a feature matrix with samples in columns and the detected mass features in rows. Features with missing values in more than 20% of the samples were removed. The remaining missing values were replaced by half of the minimum value of the respective column. Measurements of technical replicates were merged by taking the mean. Log2-transformation was applied to the resulting metabolite intensity values.

#### Metabolite / lipid extraction for LUMC measurements

Cells were extracted using a biphasic methanol/chloroform/water protocol. Briefly, 500 μL of pre-cooled methanol was added to each well, and cells were scraped to detach and transferred into 2 mL Eppendorf tubes. After all samples were collected, tubes were vortexed for 30 s and placed on wet ice. Subsequently, 250 μL of pre-cooled chloroform was added, vortexed for 30 s, and kept on ice. An additional 250 μL of chloroform and 400 μL of LC-grade water were then added, followed by vortexing for 40 s. Samples were incubated on wet ice for 10 min to allow phase separation and centrifuged at 4000 RCF for 10 min. at 4 °C. From the resulting biphasic mixture, 650 μL of the upper polar phase was collected into glass vials for untargeted metabolomics, while 500 μL of the lower non-polar phase was collected, split into two vials (90% for targeted lipid analysis and 10% for untargeted lipidomics). All extracts were dried and stored at −80 °C until analysis.

#### Metabolomics (LUMC)

Dried samples were thawed at room temperature and reconstituted in 100 μL of water, vortexed briefly, and sonicated for 1 min. Samples were transferred to inserts, and a pooled quality control sample was prepared. Metabolites were measured in liquid chromatography tandem mass spectrometry (LC-MS/MS) system, using an in-house metabolite library [54]. Briefly, 10 µL of the extracts were injected in a Shimadzu Nexera X2 (consisting of two LC30AD pumps, a SIL30AC autosampler, a CTO20AC column oven and a CBM20A controller) (Shimadzu, the Netherlands). The mobile phases, i.e. water (eluent A) and methanol (eluent B), both containing 0.1% formic acid, were delivered at 400 µL/min with the following composition: 0% B at 0 min, 0% B at 1.5 min, 97% B at 9.9 min, 97% B at 12.9 min, 0% B at 13.0 min and 0% B at 13.8 min. The column used was a Synergi Hydro-RP, 2.5 µm, 100 × 2 mm with a Phenomenex SecurityGuard Ultra C8, 2.7 µm, 5 × 2.1 mm guard column cartridge, both were kept at 40°C. The MS was a Sciex TripleTOF 6600 (AB Sciex Netherlands B.V., The Netherlands) operated in positive and negative ESI mode, with the following conditions: ion source gas 1 50 psi, ion source gas 2 50 psi, curtain gas 30 psi, temperature 500 °C, acquisition range m/z 75–650, ion spray voltage 5500 V (ESI +) and −4500 V (ESI-), and declustering potential 80 V (ESI +) and −80 V (ESI-). An information dependent acquisition (IDA) method was used to identify the different metabolites, with the following conditions for MS analysis: collision energy ± 10, acquisition time 80 ms and for MS/MS analysis: collision energy ± 30, collision energy spread 15, ion release delay 30, ion release width 15 and acquisition time 40 ms. The IDA switching criteria was to exclude isotopes within 4 Da for a maximum of 18 candidate ions to monitor per cycle. MS-DIAL (v5.1.230912), was used to align the data and annotate the different metabolites matching accurate mass, retention time and, in most cases, the MS/MS fragmentation pattern against the authentic chemical standards (our in-house metabolite database). Metabolites with the peak area’s RSD below 30% in the QC samples and with sample-to-blank ratio above 5 for at least 80% of the samples within the experimental groups were considered for further data analysis. Peak intensities were normalized by dividing by the total peak area of each sample.

Features with missing values in more than 20% of the samples were removed. The remaining missing values were replaced half of the minimum value of the respective column. Measurements of technical replicates were merged by taking the mean. Log2-transformation was applied on the resulting metabolite intensity values.

#### Untargeted lipidomics

Dried samples were thawed at room temperature, reconstituted in 50 μL isopropanol, vortexed briefly, and sonicated for 5 min. Subsequently, 50 μL of water was added, followed by brief vortexing and sonication for 5 min. Samples were transferred to inserts, and a pooled quality control sample was prepared.

Lipidomic analysis of the lipid extracts was performed using a LC-MS/MS based lipid profiling method [55]. A Shimadzu Nexera X2 (consisting of two LC30AD pumps, a SIL30AC autosampler, a CTO20AC column oven and a CBM20A controller) (Shimadzu, ‘s Hertogenbosch, The Netherlands) was used to deliver a gradient of water:acetonitrile 80:20 (eluent A) and water:2-propanol:acetonitrile 1:90:9 (eluent B). Both eluents contained 5 mM ammonium formate and 0.05% formic acid. The applied gradient, with a column flow of 300 µL/min, was as follows: 0 min 40% B, 10 min 100% B, 12 min 100% B. A Phenomenex Kinetex C18, 2.7 µm particles, 50 × 2.1 mm (Phenomenex, Utrecht, The Netherlands) was used as column with a Phenomenex SecurityGuard Ultra C8, 2.7 µm, 5 × 2.1 mm cartridge (Phenomenex, Utrecht, The Netherlands) as guard column. The column was kept at 50 °C. The injection volume was 10 µL.

The MS was a Sciex TripleTOF 6600 (AB Sciex Netherlands B.V., Nieuwerkerk aan den Ijssel, The Netherlands) operated in positive (ESI+) and negative (ESI-) ESI mode, with the following conditions: ion source gas 1 45 psi, ion source gas 2 50 psi, curtain gas 35 psi, temperature 350°C, acquisition range *m/*z 100-1800, ion spray Voltage 5500 V (ESI+) and −4500 V (ESI-), declustering potential 80 V (ESI+) and −80 V (ESI-). An information dependent acquisition (IDA) method was used to identify lipids, with the following conditions for MS analysis: collision energy ±10, acquisition time 250 ms and for MS/MS analysis: collision energy ±45, collision energy spread 25, ion release delay 30, ion release width 14, acquisition time 40 ms. The IDA switching criteria were set as follows: for ions greater than *m/z* 300, which exceed 200 cps, exclude former target for 2 s, exclude isotopes within 1.5 Da, max. candidate ions 20. Peak intensities were normalized by dividing the total peak area of each sample.

MS-DIAL (v5.1.230912), with lipid database version Msp20230821112155, was used to align the data and identify the different lipids [56–58]. Lipids with the peak area’s RSD below 30% in the QC samples and with sample-to-blank ratio above 5 for at least 80% of the samples within the experimental groups were considered for further data analysis.

Features with missing values in more than 20% of the samples were removed. The remaining missing values were replaced by half of the minimum value of the respective column. Measurements of technical replicates were merged by taking the mean. Log2-transformation was applied to the resulting metabolite intensity values.

#### Targeted lipidomics

Comprehensive targeted lipidomics was accomplished using a flow-injection assay based on lipid class separation by differential mobility spectroscopy and selective multiple reaction monitoring (MRM) per lipid species (Lipidyzer platform). A very detailed description of lipid extraction, software and the quantitative nature of the approach can be found elsewhere [59–61]. In short, after the addition of >60 deuterated internal standards, lipids were extracted using methyl tert-butyl ether. Organic extracts were combined and subsequently dried under a gentle stream of nitrogen and reconstituted in a running buffer. Lipids were then analyzed using flow-injection in MRM mode employing a Shimadzu Nexera series HPLC and a Sciex QTrap 5500 mass spectrometer. Peak intensities were normalized by dividing the total peak area of each sample.

For further analysis, lipidomic data were normalized to cell number; features with missing values in more than 20% of the samples were removed. The remaining missing values were imputed using half of the minimum value of the respective column. Measurements of technical replicates were merged by taking the mean. Log2-transformation was applied to the resulting metabolite intensity values.

### Sample Concordance Analyses

#### Sex-concordance analyses

We assessed the concordance between the reported sex of the cell line donors and the mapping to X and Y chromosomes in CNV-WES data (Fig S4A) and sex-specific gene expression levels (Fig S4B). Both the CNVs from the WES data and measured RNA levels of *UTY* (expressed only in males) and *XIST* (expressed only in females) revealed two inconsistencies with the reported sex. Moreover, for one biological replicate of the NANS-P2 cell line, a mixed sex-specific expression was measured, indicating a possible sample contamination.

Next, a PCA score plot of the CNV-WES data revealed similar inconsistencies (Fig S4C). The PC scores of the DM1 CRISPR-Cas9 rescue line, reportedly derived from DM1-P2, closely resembled those of DM1-P2. Likewise, the CHD2 CRISPR-Cas9 line displayed higher similarity to an alternative control line (409B) than to its reported parental line (WTC).

#### Re-alignment of WES data

CRAM files were initially converted to BAM format using samtools view (v1.21) [62]. BAM files were name-sorted and converted to paired-end FASTQ files using BEDTools (v2.30) [63] bamtofastq to ensure correct pairing of read names. Read alignment to the GRCh38 reference genome was performed with BWA-MEM (v0.7.18) [64] using default parameters.

Aligned reads were filtered to retain only properly paired reads with MAPQ ≥10 using samtools and sorted by coordinates with the Picard (v3.1.1) SortSam module. Duplicate reads were identified and marked with Picard MarkDuplicates, and resulting BAM files were indexed with samtools index. Base quality score recalibration (BQSR) was performed using the GATK (v4.0.10.0) BaseRecalibrator and ApplyBQSR modules, using the 1000 Genomes Omni 2.5 variant set (hg38) as known sites. Three sets of BAM files (whole-exome sequencing, RNA-seq, and EM methylation data) were analyzed. Variant calling was restricted to a targeted panel of 50 SNPs defined in Yousefi *et al* [65]. All analyses were performed relative to the GRCh38 reference genome (GCA_000001405.15_GRCh38_no_alt_plus_hs38d1_analysis_set.fna).

#### Variant Calling

Genotypes at the targeted SNP sites were obtained using bcftools (v1.21) [66] mpileup followed by bcftools call. Genotypes were called using the multiallelic calling model (-m), producing compressed VCF outputs.

Initial VCF files were processed with vcftools (v0.1.16) [67] with a minimum depth quality threshold filter of four enforced (minDP 4). The filtered VCF files were then sorted and indexed with bcftools sort and tabix, ensuring compatibility with downstream multi-sample merging and comparison. Filtered VCF files from all three sample sets were merged using vcf-merge to create a unified multi-sample VCF file.

#### SNP-concordance Analysis

To assess genetic concordance across the three omics technology datasets, bcftools gtcheck was applied, which generated genotype concordance rates for each pairwise sample comparison. Through the SNP panel as identified by Yousefi et al. [65], we were able to construct a heatmap of all pairwise concordance scores across the samples (Fig S4D). Based on these SNP-concordances, we confirmed our previous assumptions regarding sample mix-ups. Consequently, we re-labeled the CRISPR-Cas9 edited lines for the correct parental lines and the biological replicate 3 of the NANS-P2 was excluded from the complete dataset.

### Pairwise Correlations

Pairwise correlations were computed for all possible feature pairs within each dataset and group. The groups were categorized based on experimental levels, including technical replicates, biological replicates, clones, and disease groups. Only unique feature pairs were included in the correlation analysis. Pearson correlations were calculated using the built-in cor() function in R, and the results were squared to express correlations as R-squared values.

### Principal Component Analysis

Principal Component Analysis (PCA) was conducted for each individual dataset using the standard prcomp() function in R. Sample scores for the first 10 principal components were assessed for associations with phenotypic variables using the Kruskal-Wallis test. Additionally, correlations between principal component scores and continuous phenotypic variables were evaluated using pairwise Pearson correlation. All p-values were adjusted for multiple testing using the Benjamini-Hochberg false discovery rate (FDR) correction.

### Linear models

Robust linear models were implemented using the *MASS* R package (v7.3). For each omics feature, linear models were run both with and without covariates, with the observation unit included as a covariate where applicable. All resulting p-values were corrected using FDR.

Moreover, we ran mixed linear models using the *lmerTest* R package (v3.13). These models included patient ID and biological replicate number as random effects for the intercepts. The biological replicate was nested within patient ID to reflect the hierarchical structure of the study design.

### Multi-Omics Factor Analysis

The multi-omics data sets were integrating using R/Bioconductor’s MOFA2 (version 1.14.0) [20], and Python library mofapy2 (version 0.7.2). Default settings were used, through which a maximum of 6 MOFA factors were identified. The MOFA samples’ scores and feature weights on these factors were used for downstream analysis (gene set enrichment analysis). Metabolites were putatively identified using MassBank EU by comparing their m/z values and their MS/MS fragmentation spectra [68].

### Gene set enrichment analysis

Gene set enrichment analysis was performed to analyze the ranked omics features of the MOFA factor loadings, and the ranked features as identified by the nested linear mixed models. Pathway analysis was performed using multiGSEA version 1.18.0.

### DNA methylation analyses

Using R/Bioconductor’s DSS (version 2.56.0) we identified differentially methylated regions across the cell lines. Differentially methylated regions were defined as genomic regions with at least 3 CpG sites, of which more than 50% were differentially methylated on a region of minimally 50 bp.

Furthermore, EM-seq data measured at the iPSC stage and at iNeuron stage were correlated for the DM1-P2 clones. Pearson’s correlations were calculated using the set of 100,000 CpG sites that were most variable in the original data set measured in all iNeuron samples.

## Resource availability

### Lead contact

Further information and requests for resources and reagents should be directed to and will be fulfilled by the lead contact, Purva Kulkarni.

### Materials availability

The hiPSC lines used in this study are available upon request, but may require a completed materials transfer agreement.

## Data and code availability statement

### Data FAIRification using FAIR Data Station

The FAIR Data Station was [69] used to create templates for capturing metadata according to the ISA (Investigation, Study, Assay) [70] definition. Altogether, the FAIR workflow was done and recorded with FAIRDataCube [71]. The FAIR Data Station allows to convert the filled templates into linked data formats, such as RDF (Resource Description Framework) files. All metadata regarding this multi-omics data set, ranging from the phenotypic information of the patients to the sequencing and mass spectrometry instrument settings, have been captured in the ISA-files. These files have been made available through Zenodo (https://doi.org/10.5281/zenodo.19129856).

### Raw data

The raw data for each omics layer is available in the corresponding omics-specific repositories.

- Transcriptomics and epigenomics data: Accession code: EGAC50000000964
- Metabolomics and lipidomics data: https://www.ebi.ac.uk/metabolights/reviewer33033972-a22b-478e-8e1c-b094fa651f26 (*private reviewer link*)
- Proteomics data:

*For reviewer access, log in to the PRIDE website using the following details:*

Project accession: PXD076166

Token: jIxAVTGKTyl8

### Code availability

All software code that was used to pre-process and analyze the omics data sets have been made available through the X-omics Github page (github.com/Xomics/iPSC-data-analysis).

## Ethics statement

All procedures followed were in accordance with the Declaration of Helsinki revised in 2024. The institutional ethical review committee CMO Radboudumc, Nijmegen, The Netherlands has approved to conduct studies with leftover diagnostic patient material (CMO Radboudumc dossier number: 2020-6588d). Additional informed consent was obtained from the individuals or their guardians, from whom the iPSCs were developed. Sample tissues from patients were collected under protocols METC 16-4-201 (DM1 lines), METC 2018-4525 (CHD2 lines), and CMO protocol 2020-6588 (NANS-CDG lines).

## Acknowledgements

Sequencing was performed by the SNP&SEQ Technology Platform in Uppsala. The facility is part of the National Genomics Infrastructure (NGI) Sweden and Science for Life Laboratory. The SNP&SEQ Platform is also supported by the Swedish Research Council and the Knut and Alice Wallenberg Foundation. We acknowledge the Utrecht Sequencing Facility (USEQ) for providing sequencing service and data. USEQ is subsidized by the University Medical Center Utrecht and The Netherlands X-omics Initiative (NWO project 184.034.019). Proteomics measurements were performed by the Radboud Technology Center for Mass Spectrometry supported by the Netherlands X-omics Initiative partially funded by NWO (project 184.034.019). The study was executed within the Netherlands X-omics initiative funded by the Dutch Research Council (NWO project 184.034.019), and co-funded by SIMPATHIC (European Union’s Horizon Europe research and innovation program with grant agreement No 101080249), EJP-RD (European Union’s Horizon2020 program with grant agreement 825575), and EATRIS-Plus (European Union’s Horizon2020 program with grant agreement 871096). Finally, we acknowledge Dr. Rico Derks and Dr. Yassene Mohammed for their guidance on the metabolomics and lipidomics data sets.

## Author contributions

P.A.C.t’H., A.v.G, P.K. and C.d.V. conceptualized the study. L.R., E.L. and R. M. performed the experiments to generate the iPSC neuronal models, who were supervised by H.v.B., N.N.K. and D.J.L. Formal data analysis was performed by C.d.V., C.D. and L.O’G. Data analysis code used for the analysis was developed by C.d.V. and L.O’G. Omics data FAIRification was performed by J.H. and F.B. Patient samples were provided by C.D.M.K, C.G.F., J.V. and N.N.K. C.d.V. wrote the original draft of this manuscript and was responsible for the visualization of the data. All authors reviewed the manuscript before submission. The project was supervised by P.A.C.t’H., A.v.G and P.K. Funding was acquired by P.A.C.t’H. and A.v.G.

## Supplemental information

1. Supplementary Figures.docx
2. Supplementary Table 1.xlsx - Patient cell line related information
3. Supplementary Table 2.xlsx - MOFA feature weights for omics layers
4. Supplementary Table 3.xlsx - Enriched pathways across KEGG, Reactome and WikiPathways
5. Supplementary Table 4.xlsx - Enriched lipids across MOFA factors
6. Supplementary Table 5.xlsx - Enriched pathways using MOFA 2 feature weights
7. Supplementary Table 6 All_diseases_vsControls.xlsx - List of significant associations all diseases vs Controls
8. Supplementary Table 7 DM1_P2_P3_vsControls.xlsx - List of significant associations DM1 (P2 and P3) vs Controls
9. Supplementary Table 8 DM1vsControls.xlsx - List of significant associations DM1 vs Controls
10. Supplementary Table 9 CHD2vsControls.xlsx - List of significant associations CHD2 vs Controls
11. Supplementary Table 10 NANsvsControls.xlsx - List of significant associations NANs vs Controls
12. Supplementary Table 11 Sex.xlsx - List of significant associations with Sex
13. Supplementary Table 12 All_diseases_vsControls_CORRECTED.xlsx - List of significant associations all diseases vs Controls after correction
14. Supplementary Table 13 DM1_P2_P3_vsControls_CORRECTED.xlsx - List of significant associations DM1 (P2 and P3) diseases vs Controls after correction
15. Supplementary Table 14 DM1vsControls_CORRECTED.xlsx - List of significant associations DM1 vs Controls after correction
16. Supplementary Table 15 CHD2vsControls_CORRECTED.xlsx - List of significant associations CHD2 vs Controls after correction
17. Supplementary Table 16 NANsvsControls_CORRECTED.xlsx - List of significant associations NANs vs Controls after correction
18. Supplementary Table 17 Sex_CORRECTED.xlsx - List of significant associations Sex vs Controls after correction
19. Supplementary Table 18.xlsx - Differentially methylated regions after comparing DM1-P2-C2 with the other clones from the same patient

---

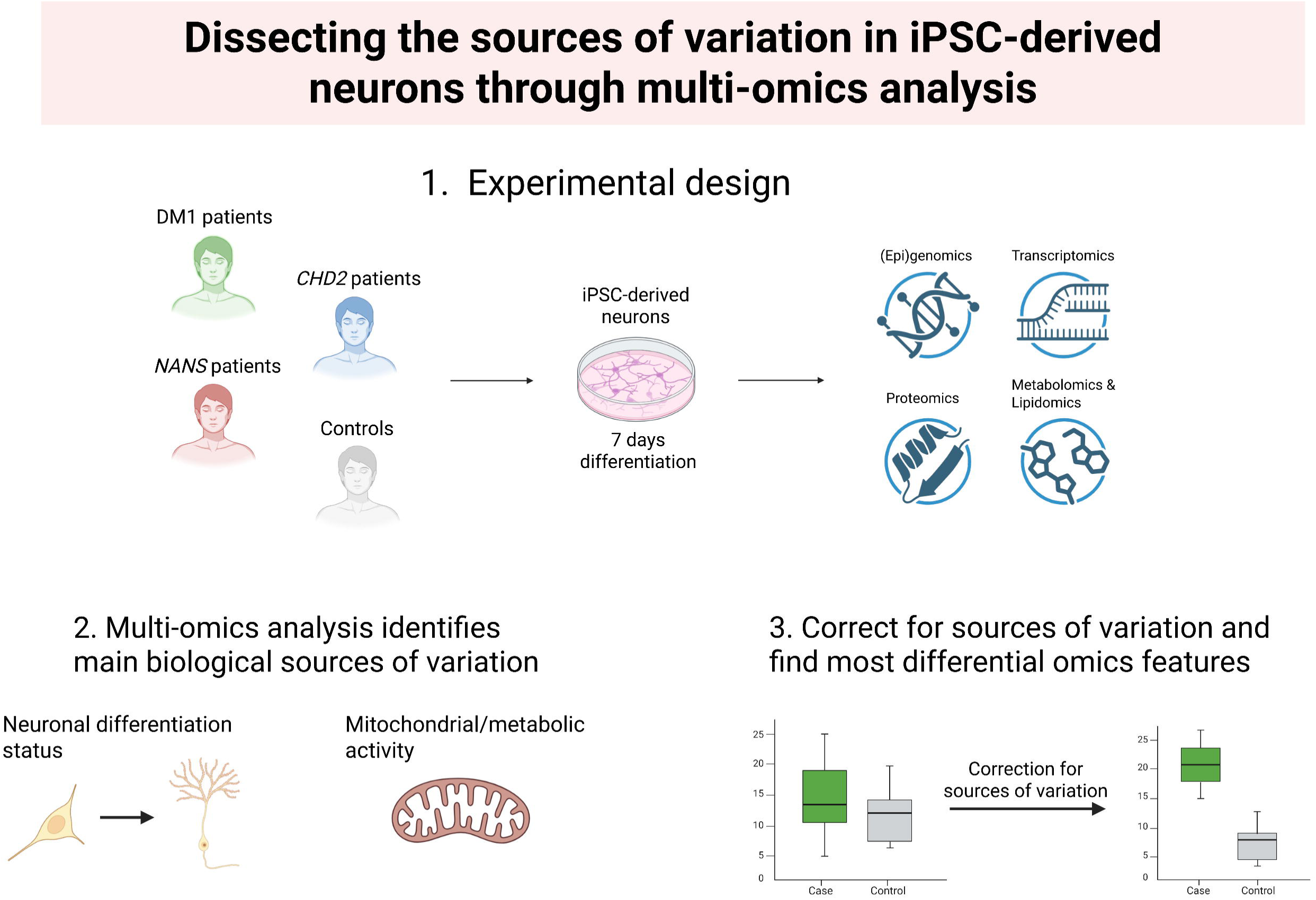

## References

[1] Moradi S, Mahdizadeh H, Šarić T, Kim J, Harati J, Shahsavarani H, et al. Research and therapy with induced pluripotent stem cells (iPSCs): Social, legal, and ethical considerations. Stem Cell Res Ther 2019;10. 10.1186/s13287-019-1455-y.

[2] Dhaiban S, Chandran S, Noshi M, Sajini AA. Clinical translation of human iPSC technologies: advances, safety concerns, and future directions. Front Cell Dev Biol 2025;13. 10.3389/fcell.2025.1627149.

[3] Kirkeby A, Main H, Carpenter M. Pluripotent stem-cell-derived therapies in clinical trial: A 2025 update. Cell Stem Cell 2025;32:10–37. 10.1016/j.stem.2024.12.005.

[4] Woodard CM, Campos BA, Kuo SH, Nirenberg MJ, Nestor MW, Zimmer M, et al. IPSC-derived dopamine neurons reveal differences between monozygotic twins discordant for parkinson’s disease. Cell Rep 2014;9:1173–82. 10.1016/j.celrep.2014.10.023.

[5] Sun X, Song J, Huang H, Chen H, Qian K. Modeling hallmark pathology using motor neurons derived from the family and sporadic amyotrophic lateral sclerosis patient-specific iPS cells. Stem Cell Res Ther 2018;9. 10.1186/s13287-018-1048-1.

[6] Kyttälä A, Moraghebi R, Valensisi C, Kettunen J, Andrus C, Pasumarthy KK, et al. Genetic Variability Overrides the Impact of Parental Cell Type and Determines iPSC Differentiation Potential. Stem Cell Reports 2016;6:200–12. 10.1016/j.stemcr.2015.12.009.

[7] Kilpinen H, Goncalves A, Leha A, Afzal V, Alasoo K, Ashford S, et al. Common genetic variation drives molecular heterogeneity in human iPSCs. Nature 2017;546:370–5. 10.1038/nature22403.

[8] Saporta MA, Grskovic M, Dimos JT. Induced pluripotent stem cells in the study of neurological diseases. Stem Cell Res Ther 2011;2:37. 10.1186/scrt78.

[9] Volpato V, Webber C. Addressing variability in iPSC-derived models of human disease: Guidelines to promote reproducibility. DMM Disease Models and Mechanisms 2020;13. 10.1242/dmm.042317.

[10] Schwartzentruber J, Foskolou S, Kilpinen H, Rodrigues J, Alasoo K, Knights AJ, et al. Molecular and functional variation in iPSC-derived sensory neurons. Nat Genet 2018;50:54–61. 10.1038/s41588-017-0005-8.

[11] Klein Gunnewiek TM, Verboven AHA, Pelgrim I, Hogeweg M, Schoenmaker C, Renkema H, et al. Sonlicromanol improves neuronal network dysfunction and transcriptome changes linked to m.3243A>G heteroplasmy in iPSC-derived neurons. Stem Cell Reports 2021;16:2197–212. 10.1016/j.stemcr.2021.07.002.

[12] Carcamo-Orive I, Hoffman GE, Cundiff P, Beckmann ND, D’Souza SL, Knowles JW, et al. Analysis of Transcriptional Variability in a Large Human iPSC Library Reveals Genetic and Non-genetic Determinants of Heterogeneity. Cell Stem Cell 2017;20:518–532.e9. 10.1016/j.stem.2016.11.005.

[13] Panopoulos AD, Smith EN, Arias AD, Shepard PJ, Hishida Y, Modesto V, et al. Aberrant DNA Methylation in Human iPSCs Associates with MYC-Binding Motifs in a Clone-Specific Manner Independent of Genetics. Cell Stem Cell 2017;20:505–517.e6. 10.1016/j.stem.2017.03.010.

[14] Yun J, So J, Jeong S, Jang J, Han S, Jeon J, et al. Transcriptome and epigenome dynamics of the clonal heterogeneity of human induced pluripotent stem cells for cardiac differentiation. Cellular and Molecular Life Sciences 2025;82. 10.1007/s00018-024-05493-9.

[15] David Brook J, Mccurrach ME, Harley HG, Buckler AJ, Church D, Aburatani H, et al. Molecular Basis of Myotonic Dystrophy: Expansion of a Trinucleotide (CTG) Repeat at the 3’ End of a Transcript Encoding a Protein Kinase Family Member. vol. 66. 1992.

[16] Hoekman TD, Rahm L, Albert S, Wansink DG, van Bokhoven H, Raaijmakers RHL. Generation of iPSC lines from myotonic dystrophy type 1 patients with varying CTG repeat lengths. Stem Cell Res 2026;94:104013. 10.1016/j.scr.2026.104013.

17. Adam MP, Bick S, Mirzaa GM. CHD2-Related Neurodevelopmental Disorders. 1993.

[18] Van Karnebeek CDM, Bonafé L, Wen XY, Tarailo-Graovac M, Balzano S, Royer-Bertrand B, et al. NANS-mediated synthesis of sialic acid is required for brain and skeletal development. Nat Genet 2016;48:777–84. 10.1038/ng.3578.

[19] Gu J, Stevens M, Xing X, Li D, Zhang B, Payton JE, et al. Mapping of variable DNA methylation across multiple cell types defines a dynamic regulatory landscape of the human genome. G3: Genes, Genomes, Genetics 2016;6:973–86. 10.1534/g3.115.025437.

[20] Argelaguet R, Arnol D, Bredikhin D, Deloro Y, Velten B, Marioni JC, et al. MOFA+: A statistical framework for comprehensive integration of multi-modal single-cell data. Genome Biol 2020;21. 10.1186/s13059-020-02015-1.

[21] Wishart DS, Guo AC, Oler E, Wang F, Anjum A, Peters H, et al. HMDB 5.0: The Human Metabolome Database for 2022. Nucleic Acids Res 2022;50:D622–31. 10.1093/nar/gkab1062.

[22] Orlicka-Płocka M, Gurda-Wozna D, Fedoruk-Wyszomirska A, Wyszko E. Circumventing the Crabtree effect: forcing oxidative phosphorylation (OXPHOS) via galactose medium increases sensitivity of HepG2 cells to the purine derivative kinetin riboside. Apoptosis 2020;25:835–52. 10.1007/s10495-020-01637-x.

[23] Ye J, Palm W, Peng M, King B, Lindsten T, Li MO, et al. GCN2 sustains mTORC1 suppression upon amino acid deprivation by inducing Sestrin2. Genes Dev 2015;29:2331–6. 10.1101/gad.269324.115.

[24] Ye J, Kumanova M, Hart LS, Sloane K, Zhang H, De Panis DN, et al. The GCN2-ATF4 pathway is critical for tumour cell survival and proliferation in response to nutrient deprivation. EMBO Journal 2010;29:2082–96. 10.1038/emboj.2010.81.

[25] Zhang D, Hua Z, Li Z. The role of glutamate and glutamine metabolism and related transporters in nerve cells. CNS Neurosci Ther 2024;30. 10.1111/cns.14617.

[26] Alfarsi LH, Ansari R El, Erkan B, Fakroun A, Craze ML, Aleskandarany MA, et al. SLC1A5 is a key regulator of glutamine metabolism and a prognostic marker for aggressive luminal breast cancer. Sci Rep 2025;15. 10.1038/s41598-025-87292-1.

[27] Fahy E, Subramaniam S, Brown HA, Glass CK, Merrill AH, Murphy RC, et al. A comprehensive classification system for lipids. J Lipid Res 2005;46:839–61. 10.1194/jlr.E400004-JLR200.

[28] Böttinger L, Horvath SE, Kleinschroth T, Hunte C, Daum G, Pfanner N, et al. Phosphatidylethanolamine and cardiolipin differentially affect the stability of mitochondrial respiratory chain supercomplexes. J Mol Biol 2012;423:677–86. 10.1016/j.jmb.2012.09.001.

[29] Nagarajan SR, Butler LM, Hoy AJ. The diversity and breadth of cancer cell fatty acid metabolism. Cancer Metab 2021;9. 10.1186/s40170-020-00237-2.

[30] Odenkirk MT, Jostes HC, Francis KR, Baker ES. Lipidomics reveals cell specific changes during pluripotent differentiation to neural and mesodermal lineages. Mol Omics 2025;21:259–69. 10.1039/d4mo00261j.

[31] Luijsterburg MS, de Krijger I, Wiegant WW, Shah RG, Smeenk G, de Groot AJL, et al. PARP1 Links CHD2-Mediated Chromatin Expansion and H3.3 Deposition to DNA Repair by Non-homologous End-Joining. Mol Cell 2016;61:547–62. 10.1016/j.molcel.2016.01.019.

[32] Steinbach P, Gläser D, Vogel W, Wolf M, Schwemmle S. The DMPK Gene of Severely Affected Myotonic Dystrophy Patients Is Hypermethylated Proximal to the Largely Expanded CTG Repeat. vol. 62. 1998.

[33] de Boni L, Gasparoni G, Haubenreich C, Tierling S, Schmitt I, Peitz M, et al. DNA methylation alterations in iPSC- and hESC-derived neurons: Potential implications for neurological disease modeling. Clin Epigenetics 2018;10. 10.1186/s13148-018-0440-0.

[34] Takahashi K, Yamanaka S. A decade of transcription factor-mediated reprogramming to pluripotency. Nat Rev Mol Cell Biol 2016;17:183–93. 10.1038/nrm.2016.8.

[35] Wu H, Wang C, Wu Z. A new shrinkage estimator for dispersion improves differential expression detection in RNA-seq data. Biostatistics 2013;14:232–43. 10.1093/biostatistics/kxs033.

[36] Wang Y, Herzig G, Molano C, Liu A. Differential expression of the Tmem132 family genes in the developing mouse nervous system. Gene Expression Patterns 2022;45. 10.1016/j.gep.2022.119257.

[37] Wang X, Jiang W, Luo S, Yang X, Wang C, Wang B, et al. The C. elegans homolog of human panic-disorder risk gene TMEM132D orchestrates neuronal morphogenesis through the WAVE-regulatory complex. Mol Brain 2021;14. 10.1186/s13041-021-00767-w.

[38] Lee G Bin, Mazli WNA binti, Hao L. Multiomics Evaluation of Human iPSCs and iPSC-Derived Neurons. J Proteome Res 2023. 10.1021/acs.jproteome.3c00790.

[39] Beekhuis-Hoekstra SD, Watanabe K, Werme J, de Leeuw CA, Paliukhovich I, Li KW, et al. Systematic assessment of variability in the proteome of iPSC derivatives. Stem Cell Res 2021;56. 10.1016/j.scr.2021.102512.

[40] Beekhuis-Hoekstra SD, Watanabe K, Werme J, de Leeuw CA, Paliukhovich I, Li KW, et al. Systematic assessment of variability in the proteome of iPSC derivatives. Stem Cell Res 2021;56. 10.1016/j.scr.2021.102512.

[41] Takahashi K, Yamanaka S. A decade of transcription factor-mediated reprogramming to pluripotency. Nat Rev Mol Cell Biol 2016;17:183–93. 10.1038/nrm.2016.8.

[42] Kim K, Doi A, Wen B, Ng K, Zhao R, Cahan P, et al. Epigenetic memory in induced pluripotent stem cells. Nature 2010;467:285–90. 10.1038/nature09342.

[43] Abyzov A, Tomasini L, Zhou B, Vasmatzis N, Coppola G, Amenduni M, et al. One thousand somatic SNVs per skin fibroblast cell set baseline of mosaic mutational load with patterns that suggest proliferative origin. Genome Res 2017;27:512–23. 10.1101/gr.215517.116.

[44] Moullan N, Mouchiroud L, Wang X, Ryu D, Williams EG, Mottis A, et al. Tetracyclines disturb mitochondrial function across eukaryotic models: A call for caution in biomedical research. Cell Rep 2015;10:1681–91. 10.1016/j.celrep.2015.02.034.

[45] Su C, Xu Z, Shan X, Cai B, Zhao H, Zhang J. Cell-type-specific co-expression inference from single cell RNA-sequencing data. Nat Commun 2023;14. 10.1038/s41467-023-40503-7.

[46] Loyfer N, Magenheim J, Peretz A, Cann G, Bredno J, Klochendler A, et al. A DNA methylation atlas of normal human cell types. Nature 2023;613:355–64. 10.1038/s41586-022-05580-6.

[47] van Agtmaal EL, André LM, Willemse M, Cumming SA, van Kessel IDG, van den Broek WJAA, et al. CRISPR/Cas9-Induced (CTG⋅CAG)n Repeat Instability in the Myotonic Dystrophy Type 1 Locus: Implications for Therapeutic Genome Editing. Molecular Therapy 2017;25:24–43. 10.1016/j.ymthe.2016.10.014.

[48] Concordet JP, Haeussler M. CRISPOR: Intuitive guide selection for CRISPR/Cas9 genome editing experiments and screens. Nucleic Acids Res 2018;46:W242–5. 10.1093/nar/gky354.

[49] Frega M, Van Gestel SHC, Linda K, Van Der Raadt J, Keller J, Van Rhijn JR, et al. Rapid neuronal differentiation of induced pluripotent stem cells for measuring network activity on micro-electrode arrays. Journal of Visualized Experiments 2017;2017. 10.3791/54900.

[50] Li H, Durbin R. Fast and accurate short read alignment with Burrows-Wheeler transform. Bioinformatics 2009;25:1754–60. 10.1093/bioinformatics/btp324.

[51] Krumm N, Sudmant PH, Ko A, O’Roak BJ, Malig M, Coe BP, et al. Copy number variation detection and genotyping from exome sequence data. Genome Res 2012;22:1525–32. 10.1101/gr.138115.112.

[52] Rajagopalan R, Murrell JR, Luo M, Conlin LK. A highly sensitive and specific workflow for detecting rare copy-number variants from exome sequencing data. Genome Med 2020;12. 10.1186/s13073-020-0712-0.

[53] Hoegen B, Zammit A, Gerritsen A, Engelke UFH, Castelein S, van de Vorst M, et al. Metabolomics-based screening of inborn errors of metabolism: Enhancing clinical application with a robust computational pipeline. Metabolites 2021;11. 10.3390/metabo11090568.

[54] Ciurli A, Mohammed Y, Ammon C, Derks RJE, Olivier-Jimenez D, Ducarmon QR, et al. Spatially and temporally resolved metabolome of the human oral cavity. IScience 2024;27. 10.1016/j.isci.2024.108884.

[55] Kong L, Dawkins E, Campbell F, Winkler E, Derks RJE, Giera M, et al. Photo-controlled delivery of very long chain fatty acids to cell membranes and modulation of membrane protein function. Biochim Biophys Acta Biomembr 2020;1862. 10.1016/j.bbamem.2020.183200.

[56] Tsugawa H, Cajka T, Kind T, Ma Y, Higgins B, Ikeda K, et al. MS-DIAL: data-independent MS/MS deconvolution for comprehensive metabolome analysis. Nat Methods 2015;12:523–6. 10.1038/nmeth.3393.

[57] Tsugawa H, Nakabayashi R, Mori T, Yamada Y, Takahashi M, Rai A, et al. A cheminformatics approach to characterize metabolomes in stable-isotope-labeled organisms. Nat Methods 2019;16:295–8. 10.1038/s41592-019-0358-2.

[58] Tsugawa H, Ikeda K, Takahashi M, Satoh A, Mori Y, Uchino H, et al. A lipidome atlas in MS-DIAL 4. Nat Biotechnol 2020;38:1159–63. 10.1038/s41587-020-0531-2.

[59] Ghorasaini M, Mohammed Y, Adamski J, Bettcher L, Bowden JA, Cabruja M, et al. Cross-Laboratory Standardization of Preclinical Lipidomics Using Differential Mobility Spectrometry and Multiple Reaction Monitoring. Anal Chem 2021;93:16369–78. 10.1021/acs.analchem.1c02826.

[60] Su B, Bettcher LF, Hsieh WY, Hornburg D, Pearson MJ, Blomberg N, et al. A DMS Shotgun Lipidomics Workflow Application to Facilitate High-Throughput, Comprehensive Lipidomics. J Am Soc Mass Spectrom 2021;32:2655–63. 10.1021/jasms.1c00203.

[61] Ghorasaini M, Tsezou KI, Verhoeven A, Mohammed Y, Vlachoyiannopoulos P, Mikros E, et al. Congruence and Complementarity of Differential Mobility Spectrometry and NMR Spectroscopy for Plasma Lipidomics. Metabolites 2022;12. 10.3390/metabo12111030.

[62] Li H, Handsaker B, Wysoker A, Fennell T, Ruan J, Homer N, et al. The Sequence Alignment/Map format and SAMtools. Bioinformatics 2009;25:2078–9. 10.1093/bioinformatics/btp352.

[63] Quinlan AR, Hall IM. BEDTools: A flexible suite of utilities for comparing genomic features. Bioinformatics 2010;26:841–2. 10.1093/bioinformatics/btq033.

[64] Li H. Aligning sequence reads, clone sequences and assembly contigs with BWA-MEM 2013.

[65] Yousefi S, Abbassi-Daloii T, Kraaijenbrink T, Vermaat M, Mei H, van ‘t Hof P, et al. A SNP panel for identification of DNA and RNA specimens. BMC Genomics 2018;19. 10.1186/s12864-018-4482-7.

[66] Danecek P, Bonfield JK, Liddle J, Marshall J, Ohan V, Pollard MO, et al. Twelve years of SAMtools and BCFtools. Gigascience 2021;10. 10.1093/gigascience/giab008.

[67] Danecek P, Auton A, Abecasis G, Albers CA, Banks E, DePristo MA, et al. The variant call format and VCFtools. Bioinformatics 2011;27:2156–8. 10.1093/bioinformatics/btr330.

[68] Horai H, Arita M, Kanaya S, Nihei Y, Ikeda T, Suwa K, et al. MassBank: a public repository for sharing mass spectral data for life sciences. Journal of Mass Spectrometry 2010;45:703–14. 10.1002/jms.1777.

[69] Nijsse B, Schaap PJ, Koehorst JJ. FAIR data station for lightweight metadata management and validation of omics studies. Gigascience 2023;12. 10.1093/gigascience/giad014.

[70] Rocca-Serra P, Brandizi M, Maguire E, Sklyar N, Taylor C, Begley K, et al. ISA software suite: Supporting standards-compliant experimental annotation and enabling curation at the community level. Bioinformatics, vol. 27, Oxford University Press; 2011, p. 2354–6. 10.1093/bioinformatics/btq415.

[71] Liao X, Ederveen THA, Niehues A, de Visser C, Huang J, Badmus F, et al. FAIR Data Cube, a FAIR data infrastructure for integrated multi-omics data analysis. J Biomed Semantics 2024;15:20. 10.1186/s13326-024-00321-2.

